# Temporal and spatial coordination of DNA segregation and cell division in an archaeon

**DOI:** 10.1101/2025.05.28.656325

**Authors:** Joe Parham, Valerio Sorichetti, Alice Cezanne, Baukje Hoogenberg, Sherman Foo, Yin-Wei Kuo, Eloise Mawdesley, Lydia Daniels Gatward, Jerome Boulanger, Ulrike Schulze, Andela Saric, Buzz Baum

**Affiliations:** MRC Laboratory of Molecular Biology; Cambridge, UK; Institute of Science and Technology Austria; Klosterneuberg, Austria; Kings College London; London, UK

## Abstract

Cells must coordinate DNA segregation with cytokinesis to ensure that each daughter cell inherits a complete genome. Here, we explore how DNA segregation and division are mechanistically coupled in archaeal relatives of eukaryotes, which lack CDK/Cyclins. Using live cell imaging we first describe the series of sequential changes in DNA organisation that accompany cell division in *Sulfolobus*, which computational modelling shows likely aid genome segregation. Through a perturbation analysis we identify a regulatory checkpoint which ensures that the compaction of the genome into two spatially segregated nucleoids only occurs once cells have assembly a division ring - which also defines the axis of DNA segregation. Finally, we show that DNA compaction and segregation depends, in part, on a ParA homologue, SegA, and its partner SegB, whose absence leads to bridging DNA. Taken together, these data show how regulatory checkpoints like those operating in eukaryotes aid high-fidelity division in an archaeon.

## Introduction

The regulation of the eukaryotic cell division cycle is now relatively well understood in a variety of model systems, from yeast to humans^1^. In eukaryotes, orderly cell cycle progression relies on waves of transcription^2^, proteasome-mediated protein degradation^3^, and oscillations in CDK-cyclin-dependent protein phosphorylation^4^. These controls are aided by cell cycle checkpoints, which ensure that critical events in the cell cycle are completed before the initiation of subsequent steps^5^. One of the best studied examples of this is the spindle assembly checkpoint, which ensures that spindle-dependent DNA separation and cytokinesis are not triggered until the system is in a state of readiness^6–8^. In addition, to ensure that each nascent daughter cell inherits a complete copy of the parental cell’s genome, DNA segregation and cytokinesis must be spatially coordinated so that the division plane bisects the center of the spindle. In different eukaryotes this coordination is achieved in distinct ways. In animal cells, for example, which undergo a relatively open mitosis, a signal emanating from overlapping microtubules at the center of the anaphase spindle triggers the assembly and contraction of an overlying actomyosin-based cytokinetic ring^9–11^. Later, the midbody positions the ESCRT-III machinery to bring about abscission at a site adjacent to the center of the division bridge^12^. Conversely, in budding yeast cells, which undergo a closed mitosis, in addition to the operation of a spindle checkpoint which ensures high fidelity DNA segregation^13^, DNA segregation and cell division are coordinated by a checkpoint that monitors spindle position^14^. Outside of eukaryotes, there is little evidence to support the operation of similar cycle checkpoints. While it is often assumed that this reflects the lack of clear temporal separation between DNA replication, DNA segregation and cell division in prokaryotes^15^, including in some archaea^16^, many of the closest archaeal relatives of eukaryotes, including *Sulfolobus acidocaldarius*, possess an orderly cell division cycle^17^. This involves waves of transcription^18^ and proteasome dependent protein degradation^19^. Furthermore, despite lacking close homologues of CDK-Cyclins^19^, *Sulfolobus* cells use eukaryotic-like Cdc6/Orc1 proteins to fire multiple origins of DNA replication once per cell cycle^20,21^ and, like human cells, employ composite ESCRT-III polymers to trigger abscission at the end of each cell cycle^22–26^. This suggests the possibility that there may be deeply conserved mechanisms of cell cycle control that have yet to be revealed.

At the same time, some of the machinery *Sulfolobus* cells use to control critical events in their cell cycle are profoundly different from those operating in eukaryotes. For example, *Sulfolobus* cells use an archaeal specific protein CdvA rather than ESCRT-I and II complexes to nucleate ESCRT-III polymer formation^27–30^. In addition, while *Sulfolobus* cells possess a Rad50-like SMC protein, called Coalescin^31^, they appear to lack functional homologues of the SMC proteins Cohesin and Condensin that help drive DNA individualization in bacteria and eukaryotes^32,33^. In addition, they lack homologues of tubulin, while possessing a ParA-like protein SegA, which based on *in vitro* work has been proposed to work together with a DNA binding-partner protein, SegB, to drive genome segregation^34,35^.

Since it is not known how archaea coordinate DNA segregation and cell division, in this paper we use *Sulfolobus* as an experimentally tractable model system to explore how these different processes function together to ensure that each daughter cell inherits a full copy of the genome at the end of each cell cycle^36^. This work reveals a complex choreographed set of changes in genome organisation that accompany division. In addition, it identifies a regulatory decision point in the *Sulfolobus* cell cycle that, like the spindle assembly checkpoint in eukaryotes, acts to ensure that cells do not commit to cell division until everything is in place - implying the existence of more general rules that aid cell division across the tree of life.

### DNA segregation and cell division are temporally coordinated

To investigate the spatial and temporal coordination of DNA segregation and cell division in *Sulfolobus*, we began by using live imaging to follow wildtype cells (DSM 639) as they progressed through the cell cycle. Cell Mask Deep Red Plasma Membrane Stain and SYBR Safe were used to label the cell membrane and DNA respectively, and cells were imaged in 15 second intervals at 75°C using our upgraded Sulfoscope set up^37,38^. This revealed a dynamic series of changes in DNA localisation, compaction and segregation that accompany the progression of cells from G2 into division (Fig 1A).

**Figure 1.**
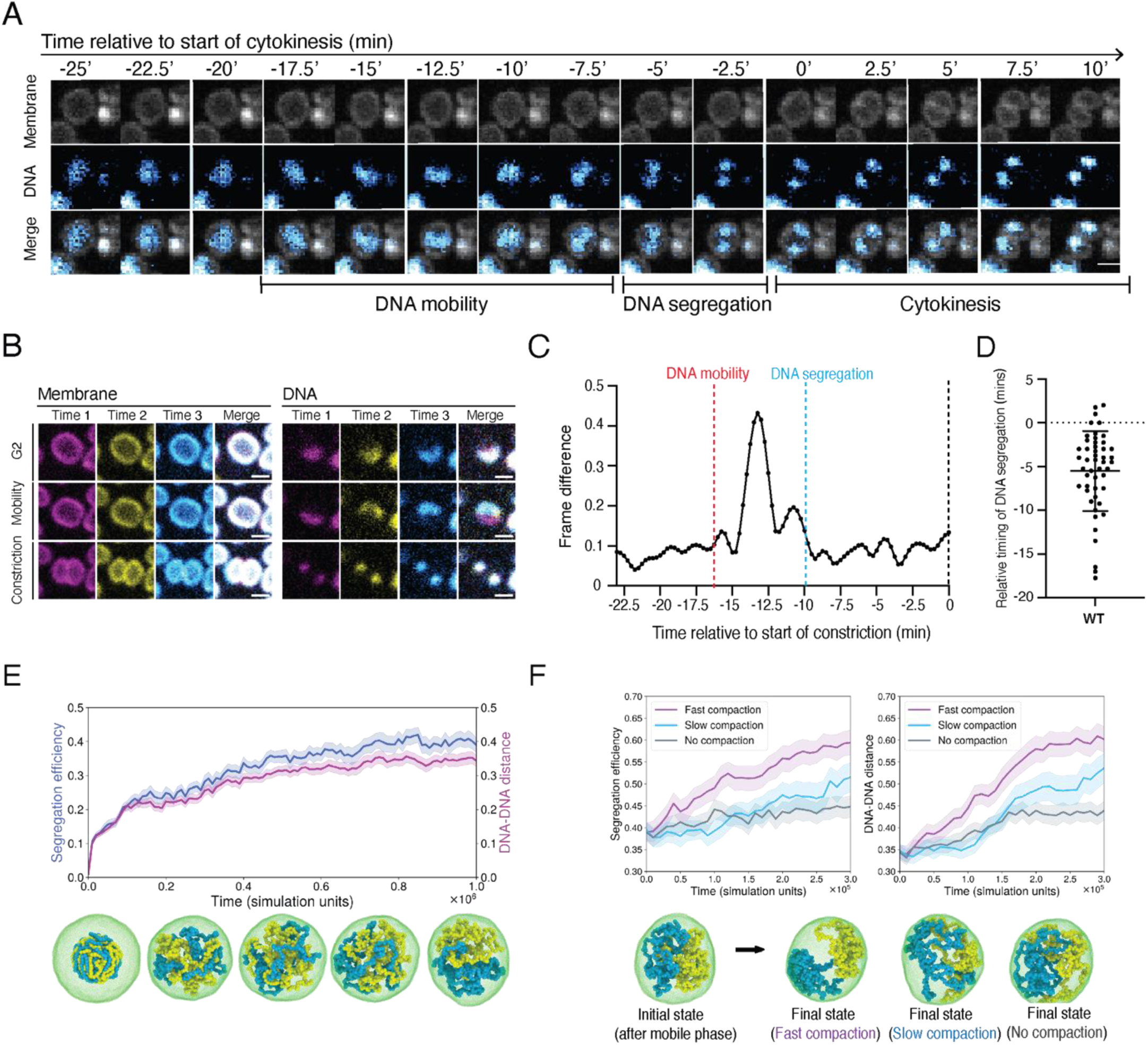
DNA segregation and cell division are temporally coordinated. (**A)** Montage of a dividing wildtype cell with membrane and DNA visualised from late G2 through DNA segregation and cell division. Scale bar 1μm. **(B)** Single slices and overlays of membrane and DNA in three frames from live imaging of a wildtype cell in three distinct phases; G2 (prior to division phase entry); DNA mobility (prior to DNA segregation) and during cell constriction, after mobility and DNA segregation. Scale bar 1μm. **(C)** An analysis of DNA motion over time in a wildtype cell made by comparing the DNA signal across frames to highlight differences. Annotations have been added by hand to highlight the discrete phases of DNA mobility, DNA segregation and the onset of cytokinesis. **(D)** Quantification of timing of DNA segregation relative to timing of the first frame of cell constriction in cytokinesis (0) in wildtype cells (n=49, N=3 biological replicates). Error bars denote mean ± standard deviation. **(E)** Segregation efficiency and normalized distance between the centers of mass of the chromosomes in simulations of cells undergoing the mobile phase (n=50; shaded area: standard error) and snapshots of one of the simulated systems for 10% DNA volume fraction. The cell membrane is coloured in green, while the two chromosomes are coloured in yellow and blue, respectively. The snapshots were taken at time intervals of 2.5×10^5^τ (simulation time units). **(F)** Segregation efficiency (left) and normalized distance between the centers of mass of the chromosomes (right) in simulations of cells undergoing fast and slow DNA compaction, compared with the no compaction control (n=50; shaded area: standard error). The data are for 10% DNA volume fraction. The snapshots represent the common initial state (left) and the final states (right) of three simulations, respectively of fast, slow and no compaction.

For most of the cell cycle, the DNA appeared stably localized to a discrete site close to the cell membrane (Figure 1A, Fig S1). However, as cells progress towards division, the DNA dissociated from the membrane - becoming diffuse and mobile in the process (Supplementary movies 1-2). After a variable length of time (average 18 ± 8.5 std, mins) (Fig S2A), this “mobile phase” ended when the DNA compacted to form two spatially separate foci. This process of nucleoid individualization was rapid, often occurring within the 15 second interval separating consecutive frames of the movie (Fig S2, Supplementary movies 1-2). Once compacted, the two nucleoids remained relatively stationary. Later, during cytokinesis, they re-associated with the membrane at opposing poles of the dumbbell-shaped doublet.

To visualize these dynamic changes in DNA organization in another way, we overlayed the DNA and membrane signals taken from the same cells at different times points in different colours (Fig 1B). Again, the DNA appeared largely stationary in G2 cells (white), highly mobile in cells preparing for division (multi-coloured), before becoming rapidly compacted to form two spatially separate stable DNA foci (white). As a more quantitative measure of DNA dynamics in cells progressing from G2 into division, we also used an image analysis pipeline (see methods) to calculate the difference in the DNA signal at each pixel over time in the subset of dividing cells that remained stationary throughout the process (Fig 1C). Consistent with the observations described above, a significant peak in DNA motion was visible in these plots prior to the onset of DNA compaction and individualization (Fig 1C), with little movement thereafter.

Importantly, the live imaging data also revealed a striking temporal correlation between the compaction of DNA into two spatially segregated nucleoids and the onset of cell division. In almost all cases, DNA segregation was complete before cytokinesis; occurring an average of 5.5 mins before the first visible cell constriction (Fig 1D). This implies strict temporal coordination between the two processes. Furthermore, DNA compaction and cell division were also coordinated in space, since the axis of DNA segregation was consistently oriented at 90° (±10.3°) with respect to the cytokinetic furrow (Fig S2). Finally, cytokinesis itself tended to be fast (∼6.6± 2.5 mins) – and was considerably less variable in duration than the mobile phase (Fig 1A, Fig S2A). Taken together, these data reveal that cell division in *Sulfolobus* is accompanied by a complex choreographed series of changes in DNA organization. In addition, they reveal a strong correlation between the axis along which the DNA is separated and compacted, and the placement of the cytokinetic furrow, implying the strict coordination of DNA segregation and division in both space and time.

As a way of assessing the likely function of the changes in DNA organization observed via live cell imaging, we developed a computational model of the process in which two circular DNA molecules (modelled as bead-spring polymers) were confined inside a vesicle, bounded by a thin membrane (Fig 1E). To begin, we used this model to analyse the role of the mobile phase in DNA segregation. Earlier theoretical studies of bacterial DNA segregation had suggested that the weak entropic repulsion between overlapping chromosomes would be sufficient to lead to their segregation^39–41^. However, this work also highlighted the importance of an elongated cell shape for efficient entropic segregation, implying that little to no segregation would occur via this mechanism in spherical cells^41^.

To take a fresh look at this problem and investigate whether entropic forces can lead to significant segregation in spherical cells, like those of *Sulfolobus,* we initiated simulations with DNA in a compact state (representing volume fractions of either 5, 10 or 20%). We then let the DNA relax and undergo Brownian motion, while allowing chromosome strands to cross in order to model the function of Topoisomerase-II. To assess the success of nucleoid segregation in simulations, we employed two different metrics, both of which tended to give similar results. The first, which we term “segregation efficiency”, was obtained using a Linear Discriminant Analysis (LDA). This quantity is equal to 0 for a perfectly mixed configuration and 1 for a perfectly segregated one. The second measure defines the distance between the centers of mass (CoM) of the two chromosomes, normalized by the vesicle radius. By analysing simulations in this way, we were able to show that entropic forces are sufficient to drive the partial segregation of chromosomes in a spherical vesicle, in a manner that is improved by lowering the DNA volume fraction^41^ (Figure 1E and Fig S3). When acting alone, however, these entropic forces were not sufficient to induce full DNA segregation (Fig 1E), since they are unable to maintain the disentangled state.

To explore whether rapid compaction of DNA observed by live imaging might facilitate DNA segregation (Fig 1), we modelled this process by introducing a weak nonspecific attraction between all polymer beads, which was inhibited in a plane at the boundary of the two nucleoids to ensure alignment of the division machinery with the axis of DNA compaction (see below). After a period of entropic separation sufficient to reach equilibrium, we then compared the impact of slow or fast DNA compaction on DNA segregation relative to a no compaction control. While fast compaction of the DNA was not sufficient to induce complete DNA segregation, it significantly improved the quantitative outcome of simulations (Fig 1F, Fig S3). Moreover, by combining an extended period of the mobile phase with fast compaction with the reassociation of the DNA with the membrane (Fig 1F, Fig S3), we were able to achieve an ∼60% segregation efficiency. These data imply that the different phases of choreographed DNA movement (DNA mobility, rapid DNA compaction, and the reassociation of DNA with the membrane that brings the cycle to a close) likely contribute to DNA segregation in *Sulfolobus*, while also making it clear that other mechanisms not included in this simple coarse-grained model are likely required to ensure that DNA segregation is robust and goes to completion.

### The ESCRT-III ring couples DNA segregation to cell division

Having explored how DNA dynamics likely contributes to DNA segregation, we next sought to determine how DNA segregation, division ring assembly, and cytokinesis are coordinated. This requires aligning the axis of DNA segregation with the plane of cytokinesis. We considered two alternative hypotheses by which this might occur: i) the machinery driving DNA segregation defines the site of the future division plane (as the spindle does in human cells), or ii) the cytokinetic machinery sets the division plane to ensure the coordination of DNA segregation and cytokinesis. To ascertain which of these hypotheses best reflects the picture in dividing *Sulfolobus* cells, it was necessary to determine the relative timing of different steps in the process of division ring assembly with respect to the observed changes in DNA organization using immunofluorescence.

To begin this analysis, we extended our previous description of the steps in the pathway of ESCRT-III ring assembly^19,24,37^ by determining the timing of CdvA expression and ring formation. Cells were fixed at 10–20-minute intervals following release from a G2 arrest, stained for CdvA and CdvB, and analysed by flow cytometry. By gating cells with a 2N DNA content, the levels of CdvA could be compared with those of CdvB as cells progressed towards division. By this measure, levels of CdvA were found to rise 10-20 minutes before levels of CdvB (Fig S4A-B). Furthermore, using confocal microscopy, ring-like CdvA structures were seen forming at mid cell prior to formation of CdvB, B1 and B2 rings (Fig 2A, Fig S4C). These data support the previously proposed idea that CdvA acts as a template for the recruitment of CdvB^27,28^.

**Figure 2.**
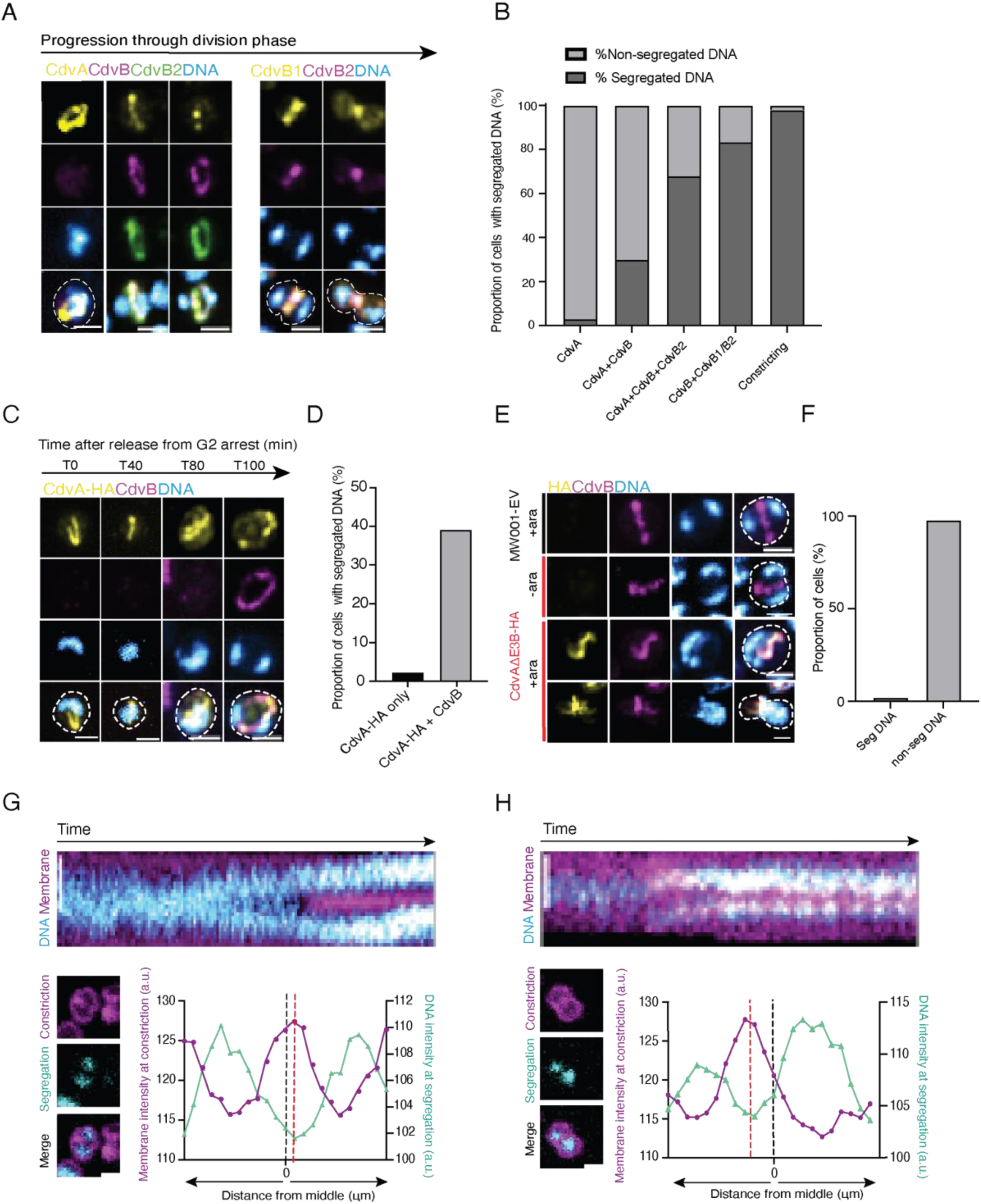
The ESCRT-III ring spatially and temporally couples DNA segregation to cell division. **(A)** Panel shows immunofluorescence images of wildtype DSM639 cells in different stages of cell division phase: cells with CdvA only rings; cells with CdvA + CdvB + CdvB2 rings; cells with CdvB + CdvB2 rings and cells in early and late constriction with just CdvB1 and CdvB2 rings. **(B)** Quantification of DNA segregation (proportion of cells with segregated vs non-segregated DNA) in wildtype cells with sequential ring compositions CdvA only (n=63), CdvA+CdvB (n=10), CdvA+CdvB+CdvB2 (n=28), CdvB+CdvB1 or CdvB2 (n=90) and constricting cells (n=65) (N=3 biological replicates). **(C)** Representative immunofluorescence images of CdvA-HA positive cells at successive time points after early induced expression in a G2 arrest. **(D)** Quantification of DNA segregation from early induction of MW001-CdvA-HA in a G2 arrest (n=66). **(E)** Representative immunofluorescence images of MW001-EV control and CdvA^ΔE3B^-HA overexpression before and after 4-hour arabinose induction. **(F)** Quantification of DNA segregation in cells expressing CdvA^ΔE3B^-HA after 4-hour arabinose induction (n=344) (N=3 biological replicates). **(G)-(H)** Kymograph of single wildtype cells through live imaging movie and single overlays of DNA signal at first frame of DNA segregation and an early frame in constriction, where the division plane is clearly defined. Plots of membrane intensity at early constriction overlayed with DNA intensity from the first frame of DNA segregation where 0 is the geometric middle of the cell. All scale bars = 1μm.

Next, to investigate the timing of DNA segregation in cells relative to ESCRT-III ring recruitment, we stained both asynchronous and synchronized populations of *Sulfolobus* cells with combinations of CdvA, CdvB, CdvB1 and CdvB2 antibodies to visualise the various stages of division. In cells with CdvA-only rings, this microscopy-based analysis revealed that the DNA formed a single diffuse mass, with only 3% of cells displaying two individualised DNA foci. In the rare cells with a composite CdvA/CdvB ring, the proportion of cells with two DNA foci rose significantly to 30%. This increased further to 68% with the recruitment of ESCRT-III homologs CdvB1 and CdvB2. Following CdvA removal, the proportion of cells with two spatial separate DNA masses reached 83%. Finally, 98% of constricting wildtype cells were found to possess two individualized DNA foci (Fig 2B, Fig S4D). These data reveal a tight temporal coupling of DNA segregation and cytokinesis. In addition, the data suggest that, in addition to its role as a template for the recruitment of the ESCRT-III division machinery, CdvA may also serve as a spatial marker that helps to define the plane of DNA separation.

Since DNA segregation follows the recruitment of CdvB to the CdvA ring, we considered several possible modes of regulation. First, the process might be triggered by some unknown target of the pre-division wave of gene expression, with some delay relative to CdvA. Alternatively, the cue for DNA segregation might depend on assembly of the CdvA ring or on formation of the ESCRT-III division ring itself. To distinguish between these different possibilities, we asked whether the premature accumulation of CdvA protein was sufficient to induce DNA segregation. To do so, we overexpressed full-length CdvA tagged with HA from an inducible arabinose promoter in G2 arrested cells. Upon release from the arrest, cells were fixed and stained at 20-minute intervals as they exited G2 and entered division. Strikingly, under these conditions, ectopically expressed CdvA was able to form linear, ring-like structures a full 80 minutes before the rise of endogenous CdvA expression in the MW001-empty vector control (Fig 2C, Fig S4E). The presence of CdvA ring-like structures, however, was not sufficient to induce DNA segregation. Instead, significant DNA segregation was only observed 100 minutes post-release, coincident with the recruitment of CdvB to the CdvA ring (Fig 2D).

Next, to determine whether DNA segregation can occur in cells that are unable to complete ring assembly, we ectopically expressed a truncated version of CdvA (hereafter named CdvA^Δ^_E3B_) that lacks the CdvB-binding E3B domain (Fig S5A)^27,28^. This was sufficient to prevent the formation of complete CdvB rings, leading to the formation of abnormal linear CdvA positive structures that likely contain both endogenous CdvA as well as the CdvAΔ_E3B_ mutant protein. While these linear structures were able to recruit a portion of the cellular pool of CdvB, the expression of CdvA^Δ^_E3B_ almost completely blocked the recruitment of CdvB1 (Fig S5B-C). As a consequence, cells expressing truncated CdvA were unable to divide, as evidenced by both the accumulation of large cells that had failed division (Fig S5D) and by the decrease in the percentage of 1N cells observed by flow cytometry (Fig S5E). At the same time, these cells expressing CdvA-HA^Δ^_E3B_ and CdvB failed to compact and individualize the two copies of their replicated genome (Fig 2E-F). These data imply that DNA segregation does not occur until after the assembly of a functional ESCRT-III division ring. Furthermore, in this experiment we observed the small number of cells expressing CdvA-HA^Δ^_E3B_ that slipped through the division arrest, which contained a single mass of DNA on one side of the partially constricted defective division ring (Fig 2F) - reinforcing the idea that DNA segregation requires the assembly of a fully functional division ring. Similarly, in live imaging experiments, cells expressing CdvA^Δ^_E3B_ remained stuck in a state with highly mobile DNA for up to 3 hours, as if unable to compact and segregate their DNA and divide (Supplementary movies 3-5).

Next, to determine how cells align the axis of DNA separation relative to their future cytokinetic furrow, we analysed live imaging data to generate a set of line profiles perpendicular to the future division plane which we could then use to compare dynamic changes in the DNA and membrane signal. These two-colour kymographs revealed that DNA segregation occurs around the future plane of membrane constriction (Fig 2G). By measuring membrane signal intensity across the whole cell during the earliest phase of constriction, we were then able to determine the position of the future cytokinetic furrow, which we could use as a landmark to compare with the position at which the gap between the new condensing nucleoids first becomes visible, a few minutes earlier. While limits in the resolution of light microscopy made it difficult to determine with high confidence whether the DNA segregates away from the geometric cell center or away from the site of the future furrow in cells that divide symmetrically (Fig 2G), in rarer asymmetric cell divisions it was clear that the DNA is cleared from a site that pre-figures the division plane rather than being aligned with the middle of the cell (Fig 2H). Taken together, these results show that maturation of the division ring through the recruitment of ESCRT-III proteins is a pre-requisite for passage through the regulatory decision point that triggers the onset of DNA segregation and cytokinesis.

### SegA and SegB regulate DNA compaction

Having shown that assembly of the ESCRT-III division ring is a pre-requisite for DNA segregation, we next wanted to investigate the machinery involved in DNA compaction and individualisation itself. As potential regulators, we turned our attention to the proteins SegA and SegB, since these proteins are expressed at the same time as ESCRT-III proteins in preparation for cell division^42^, and have been suggested to bind DNA to drive chromosome segregation in *Sulfolobus*^34,35^.

To investigate their roles in DNA segregation and division we used antibodies raised against SegA and SegB^43^ (kindly given to us by Arthur Charles-Orszag) to visualise the localization of the endogenous protein in cells in different stages of cell division (Fig 3A). In wildtype cells that had not yet undergone DNA segregation, SegB was seen localising in puncta on the DNA^42^. As the DNA became more compact, these structures tended to coalesce into two well-defined loci, which remained in place throughout DNA segregation and cytokinesis, as expected if SegB binds SegS sites as previously proposed^34,44^, and similar to those observed in other recent studies^43,44^. At the same time, many cells contained additional foci that were less obviously associated with the main nucleoid mass. We also used immunolocalization to investigate localization of SegA^43^ in a similar manner. While there was a high background, the majority of the antibody signal was detected on or close to the DNA during division. We note that Orszag et al., also proposed that there is a focus of SegA staining between well-separated nucleoids^43^. These data are consistent with the possibility that SegA and SegB might play a role in DNA compaction and segregation at both early and late stages of division (Figure 3A).

**Figure 3.**
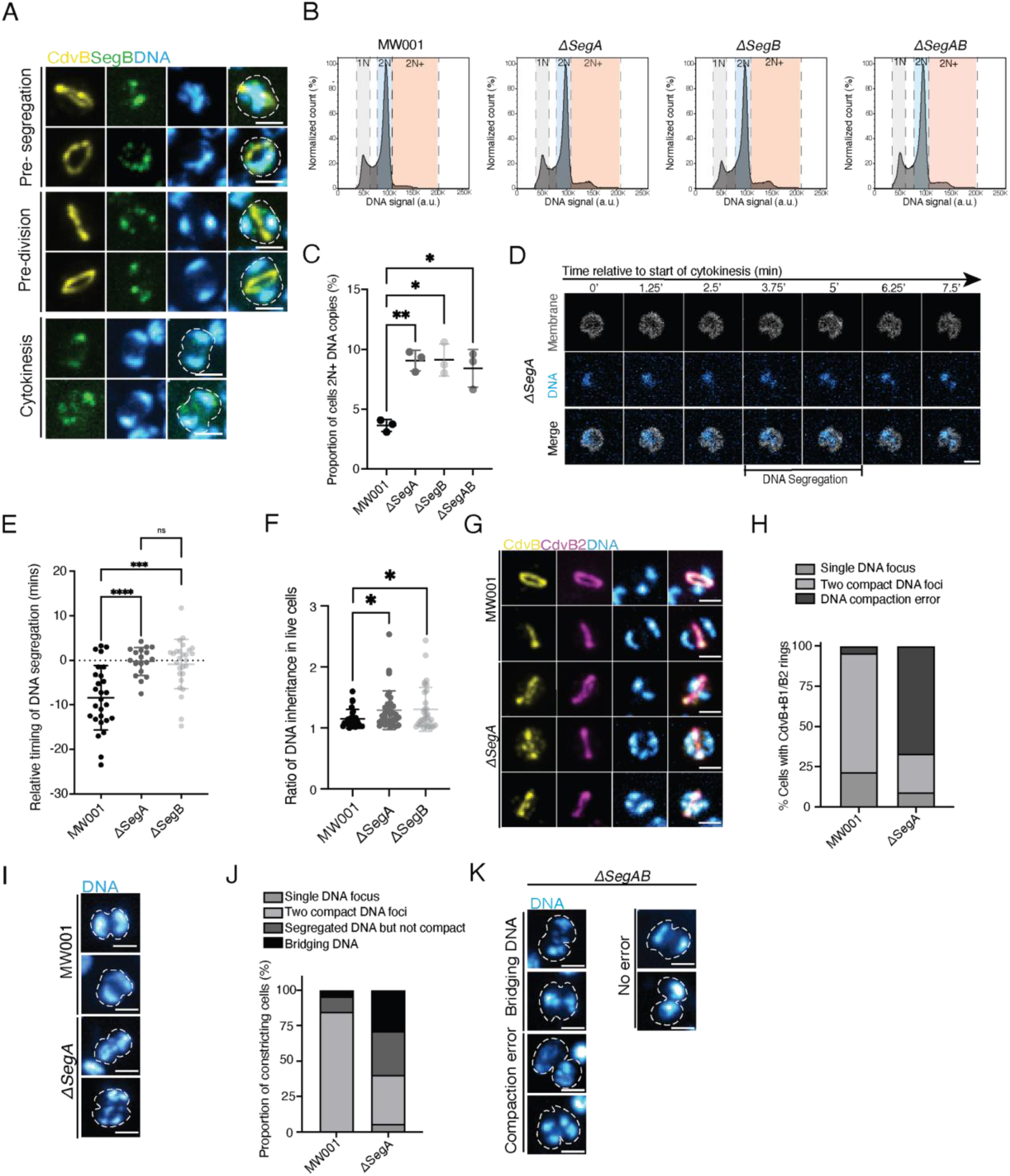
SegAB temporally coordinates DNA compaction with cell division. **(A)** Representative immunofluorescence images of control DSM639 cells stained with ESCRT-III protein CdvB (yellow), DAPI (blue) and SegB (green) in different stages of division phase; pre-DNA segregation, post-DNA segregation and cytokinesis. All scale bars = 1μm. **(B)** Representative flow cytograms of control MW001 cells and gene deletion strains *ΔSegA, ΔSegB* and *ΔSegAB* showing a slight increase in the 2N+ cell population, n=0.25X10^6^ (N=3 biological replicates). **(C)** Quantification from flow cytograms of proportion of cells in populations of MW001, *ΔSegA, ΔSegB* and *ΔSegAB* that have 2N+ copies of DNA. Significance values were derived using Welsch’s t-test (N=3 biological replicates) Error bars denote mean ± standard deviation. MW001 vs *ΔSegA, P-value =* 0.0019, MW001 vs *ΔSegB, P-*value=0.0113, MW001 vs *ΔSegAB, P-*value= 0.0250. (**D)** Montage of dividing *ΔSegA* cell from the first frame of constriction in live imaging where cells segregate their DNA asymmetrically after the onset of cytokinesis. **(E)** Quantification of timing of DNA segregation relative to timing of the first frame of cell constriction in cytokinesis (0) in control MW001 (n=27), *ΔSegA* (n=18) and *ΔSegB* (n=27) cells. *P*-values were derived by Mann-Whitney U unpaired test (N=3 biological replicates) MW001 vs *ΔSegA, P-*value=0.0001, MW001 vs Δ*segB P*-value=0.0002. **(F)** Quantification of DNA asymmetry in inheriting daughter cells from live imaging of MW001(n=26), *ΔSegA* (n=34) and *ΔSegB* (n=32). *P*-values were derived by Welch’s t-test. MW001 vs *ΔSegA, P-*value= 0.0240, MW001 vs *ΔSegB, P*- value=0.0304. Error bars denote mean ± standard deviation **(G)** Control MW001 and *ΔSegA* cells with ESCRT-III rings in division phase stained with CdvB (yellow), CdvB2 (magenta) and DAPI (blue) to show DNA compaction. **(H)** Quantification of DNA compaction (compact into two foci, compaction error, or not yet segregated) in MW001 control (n=46) and *ΔSegA* (n=33) cells that have CdvB+CdvB1/B2 composite rings (N=3 biological replicates). **(I)** MW001 control cells and *ΔSegA* cells fixed during cytokinesis and stained with DAPI (blue). **(J)** Quantification of DNA state in these constricting cells (Single DNA focus, compact into two foci, compaction error or bridging DNA)(MW001 n=46, *ΔSegA* cells n=52) (N=3 biological replicates). **(K)** Constricting Δ*segAB* cells with bridging DNA, compaction error and no error. All scale bars = 1μm.

To test this idea and determine whether SegA and SegB play an active role in DNA segregation, we generated knockout strains for SegA *(ΔSegA)* and SegB *(ΔSegB)* as well as a double knockout strain lacking both proteins *(ΔSegAB)* – all of which were viable and grow well. By flow cytometry analysis, the loss of SegA and/or SegB led to a small, but statistically significant increase in the number of cells with more than two copies of the genome relative to the control (Fig 3B-C).

To determine the origin of this modest division phenotype, we imaged *ΔSegA* and *ΔSegB* cells live (Fig 3D-F, Fig S6). In interphase, mutant cells appeared similar to the MW001 control. However, while DNA became mobile as *ΔSegA* and *ΔSegB* cells entered division, as usual, there was a profound change in the transition from mobility to DNA segregation. Most strikingly, many cells underwent DNA segregation after the onset of membrane constriction - something rarely seen in the corresponding MW001 control (Fig 3E, Fig S6). Despite this, most *ΔSegA* and *ΔSegB* cells were still able to visibly compact and separate their DNA into two foci by late stages of cytokinesis. Similarly, when the sum intensity of DNA in each daughter cell during cytokinesis was quantified, we saw only a marginal increase in the average asymmetry of DNA inheritance in mutant cells as compared to the MW001 control (Fig 3F). Taken together these data show that while SegA and SegB play an important role in the temporal coordination of DNA segregation and cell division, back-up systems appear to be able to aid DNA segregation in their absence.

To investigate whether SegA and SegB play a role in DNA compaction itself, we investigated DNA morphology at higher resolution in mutant cells that had been fixed and stained using confocal microscopy. We focused on the analysis of DNA morphology in cells possessing a full ESCRT-III ring, as determined by the presence of CdvB and CdvB1 or B2, at a time when DNA segregation would normally be complete (Figure 2). In contrast to the control, *ΔSegA, ΔSegB* and *ΔSegAB* cells frequently failed to compact their DNA into two foci (Figure 3G-H,Fig S7). Instead, the DNA was seen in bright spots that were connected by thin linear segments.

This phenotype persisted in *ΔSegA* and *ΔSegB* cells that had been fixed mid-constriction. In these cells, DNA was often seen bridging the cytokinetic furrow - a phenotype that was extremely rare in dividing MW001 control cells (Figure 3 I-J). Although these phenotypes appeared striking, a subset of dividing cells in all three deletion strains had correctly compacted and segregated DNA. Again, these data imply the existence of additional machinery that acts together with SegA and SegB to aid DNA segregation (Fig 3K,Fig S7). Furthermore, while the overexpression of the *SegAB* cassette under the control of an inducible promoter generated proteins that localized correctly to the DNA, their expression was not sufficient to induce compaction or DNA segregation (Fig S8). Taken together these results suggest that while SegA and SegB are not sufficient to induce premature DNA compaction and individualization, and are not essential for DNA segregation, they play a significant role in the temporal coupling of DNA compaction and separation with cell division in *Sulfolobus*.

## Discussion

In this study, we used *Sulfolobus acidocaldarius* as a model to characterise the series of events that ensure the correct partitioning of a single copy of the genome into two daughter cells in an archaeal relative of eukaryotes. This work reveals the existence of a regulatory decision point in the archaeal cell cycle that *Sulfolobus* cells use to coordinate DNA segregation with cytokinesis (Figure 4). This functions to achieve the same goal as the spindle checkpoint in human cells and the spindle positioning checkpoint in yeast^45,46^, to prepare cells before they commit to division and to re-entry into G1 of the following cell cycle.

**Figure 4:**
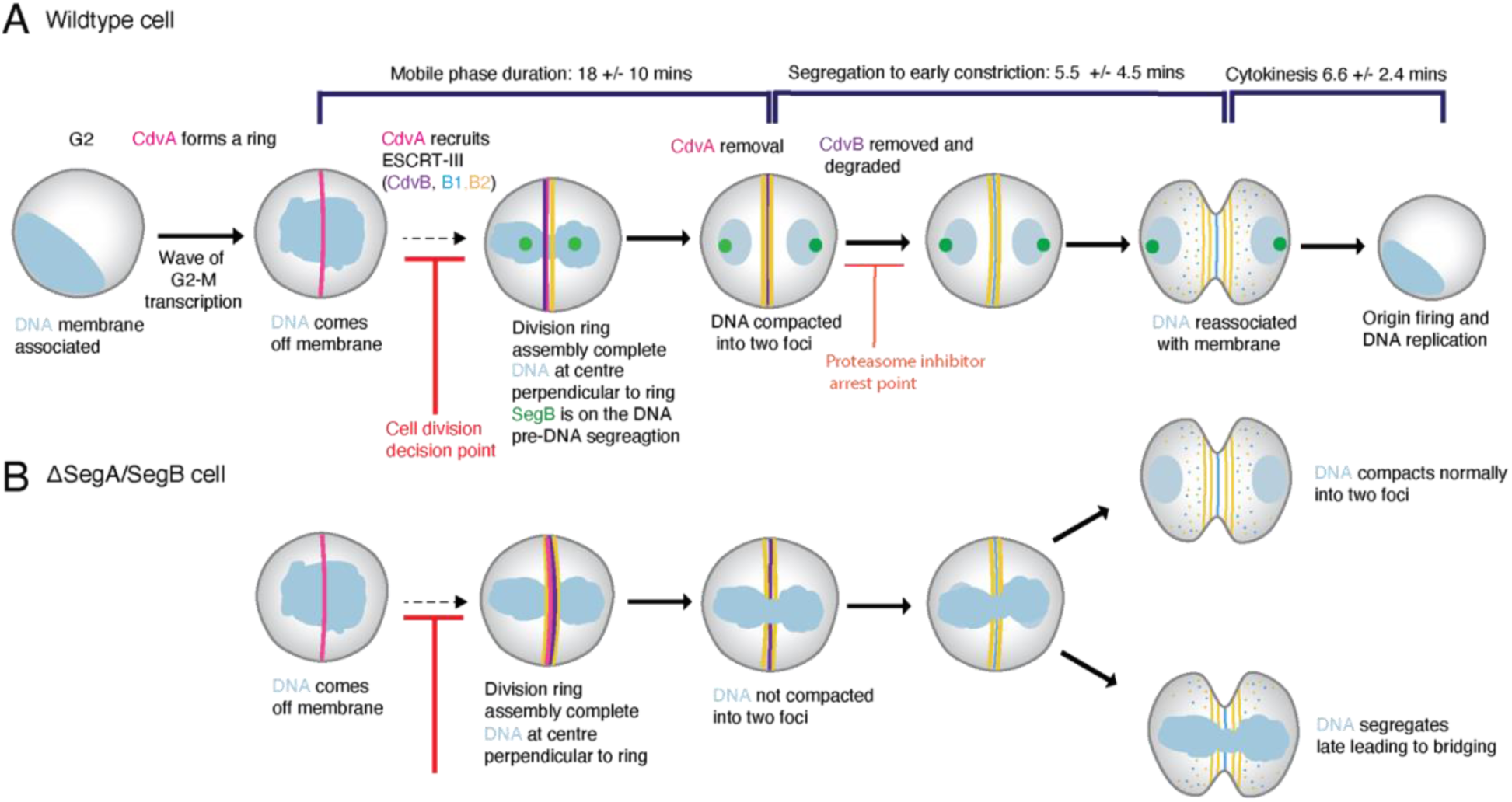
Schematic diagram of *Sulfolobus* cell division control. **(A)** Control of *Sulfolobus* cell division; cells undergo a wave of transcription at the G2-M transition where the division and segregation proteins are upregulated. CdvA forms a medial division ring before the expression of ESCRT-III but is not sufficient to induce DNA segregation. CdvA recruits ESCRT-III proteins CdvB and CdvB1 and B2 to the division ring. Perturbing full ESCRT-III ring assembly at this stage blocks DNA segregation and subsequent cell division and cell cycle progression. If cells pass through this regulatory checkpoint, the two nucleoids segregate via compaction by SegA and SegB (and other undescribed factors) before the onset of cytokinesis. CdvB is removed and degraded to allow CdvB1 and CdvB2 cells to constrict and divide before DNA relicensing and entry into G1 and S phase. **(B)** Perturbation of *Sulfolobus* cell division via SegA/SegB knockout leads to cells with late DNA segregation and compaction leading to dividing cells yet to complete DNA segregation, as seen with bridging DNA.

In *Sulfolobus* cells, as in human cells, passage through this control point is associated with the transition from a single DNA mass to two spatially separated masses. After this point, cells are committed to division and entry into G1. This implies that the action of a pre-division regulatory control point is a more ancient mechanism of cell cycle control than coupling cell cycle progression to oscillations in the levels of CDK cyclins.

As we show in this paper, the complex series of events that accompany division in *Sulfolobus* begin up to half an hour before division, when the duplicated genome loses its association with the bounding cell membrane - similar to the way DNA comes away from the nuclear envelope in a eukaryotic cell entering mitosis^47^. After membrane detachment, the duplicated genome becomes both diffuse and highly mobile. In fact, the localization of genomic DNA appears to change within the 15 seconds timeframe of our movies (Supplementary Movie1-2). While this mobile phase lasts a variable amount of time, it comes to an end with rapid DNA compaction.

The computer model we developed to test the likely role of these observed DNA dynamics in DNA segregation shows that entropic forces can help disentangle DNA polymers in spherical cells, like *Sulfolobus*, even though previous work on DNA segregation in bacteria suggested that such entropic forces depend on cells having an elongated shape^41^. Thus, entropy could play a more general role in genome segregation in cells with different shapes. In this light, the “mobile phase” of division likely facilitates the entropic de-mixing and individualization of the two copies of the *Sulfolobus* genome. In cells, this mobile phase appears to come to a sudden end as the DNA compacts. Our simulations show that this rapid phase of compaction can aid the separation of the DNA into two spatially separate masses in two ways: it locks in the effects of entropic separation to aid DNA segregation (hence the need for speed) and reduces DNA mobility. In the model, efficient DNA segregation is also facilitated via the reassociation of the DNA with the membrane at cell poles – something we observe at late stages of cytokinesis. Although this analysis suggests how the dynamic changes in DNA-membrane association (high in G2, low in division, high during cytokinesis) and DNA compaction (high in G2, low pre-DNA segregation and high pre-division) observed in cells likely contribute to DNA segregation, it is clear from the model that other ingredients need to be added to enable robust and complete DNA individualization, e.g. machinery that functions in a way similar to that proposed for Condensin^48^ to compact of loop DNA molecules in *cis*. Whether SegA and SegB act in this way, or whether this relies on other machinery is a question we aim to resolve in future work.

At the same time as these dramatic changes in DNA organization are underway, cells assemble a medial cytokinetic ring (see Figure 4). At the stage in the cell cycle when the DNA is mobile, our data from fixed cells suggests that the template protein CdvA begins forming a ring at the future site of cytokinesis. This CdvA ring then recruits the ESCRT-III homologue, CdvB (which lies downstream of CdvA in the CdvABC operon) likely through an interaction between the broken winged helix domain of CdvB and CdvA’s E3B tail^27^. This CdvAB ring in turn recruits the contractile ESCRT-III homologues CdvB1 and CdvB2, leading to the formation of a complete composite ring that, once activated, contracts to cut cells into two^19^.

Crucially, our data shows that the assembly of this mature composite division ring sets the stage for the subsequent events required to ensure a successful division in several ways (Figure 4). Thus, the ring not only recruits the division machinery in preparation for cytokinesis. It is also required to trigger the compaction of the DNA into two individualized masses, and the ring defines the axis of DNA segregation along which the two copies of the genome move apart from each other, towards the poles of the cell. Precisely how cells monitor completion of the division ring and used this information to direct DNA segregation is a fascinating question that remains to be determined. At the same time as DNA segregation is occurring, CdvB is removed from the pre-division ring by Vps4^24^, allowing for its subsequent degradation by the proteasome^19^, whose activity is also required during division for DNA replication in the following cycle^19^. Taken together, these data support the idea that there is a regulatory point in cell cycle when the construction of the composite division ring has been completed, which marks the end of the variable phase of DNA mobility, when cells change their state and become committed to division and entry into G1.

Interestingly, we saw no similar evidence to support the existence of an equivalent checkpoint that monitors ring constriction or DNA segregation itself, since DNA segregation and DNA replication appeared unperturbed in cells expressing a dominant negative Vps4^24^ and cells lacking SegA and SegB do not block progression into cytokinesis. Thus, as with the spindle checkpoint in human cells^46^, *Sulfolobus* cells seem to have invested all control in a single decision point prior to DNA segregation that must be passed for cells to progress into the next phase of the cell cycle.

While we hope that future work will identify the machinery used to assess ring completion and transmit this information to different elements of the downstream machinery, in this paper we show that SegA and SegB play a critical role in the temporal coordination of ring completion and DNA segregation. We show that these two proteins, which sit in the same operon, are required together for the timely compaction of DNA into two spatially separate masses following completion of the division ring. In their absence, cytokinesis is initiated prior to DNA segregation, leading to the accumulation of cells with bridging DNA.

Nevertheless, while SegA and SegB perform an important role in DNA compaction and segregation^43,44^, the fact that cells are still able to complete DNA segregation in their absence (something also observed in a parallel study by Charles-Orszag et al^43^) points to the existence of unknown molecular players. Given the importance of cell division for lineage survival, it is not surprising to find that the system is robust to perturbation. Other molecules involved may include a set of recently identified DNA binding proteins^49,50^. It is worth noting, however, that while almost all bacteria and eukaryotes rely on SMC-like proteins to individualize intertwined DNA strands by scanning the DNA in *cis* and generating loops (sometimes in partnership with bacterial ParA and ParB proteins), it is not clear whether a similar system operates in *Sulfolobus*. The only SMC protein thus-far identified in *Sulfolobales,* called Coalescin, is more similar to Rad50 than it is to Condensin or Cohesin^31^. Furthermore, the timing of Coalescin expression is significantly later than that of the ESCRT and Seg proteins^18^, suggesting a role in S or G2 phase is more likely than in DNA segregation or cell division. Given the need for additional systems to facilitate high fidelity division in our model, we think it likely however that there are non-SMC proteins that operate in an analogous way to aid chromosome individualization in *Sulfolobus*.

## Acknowledgements

We thank Matthew Kenneth for his assistance with live cell imaging. We thank Arthur Charles-Orszaig and Dyche Mullins for generously gifting the SegB antibody, and Sonja-Verena Albers for gifting the CdvA-HA overexpression plasmid. We thank the Light Microscopy and Flow Cytometry facilities at the MRC-LMB, and all the core staff at the MRC-LMB for their support. We thank all members of the Baum lab for helpful discussions. We would like to thank Arthur Radoux-Mergault, Magdalena Lechowska, Gautam Dey, Laura Downie and Iva Tolic for critical reading of the manuscript.

## Author contributions

These authors contributed equally: Valerio Sorichetti and Alice Cezanne J.P and B.B conceived the study. Initial observations were made by J.P., B.H. and B.B. Cell biology methods and experiments were planned and performed by J.P., B.H., Y.K., E.M. and L.D.G. with guidance from B.B. Molecular genetics were done by J.P., S.F. and Y.K. A.C. carried out the bulk of the live imaging experiments. Software for image analysis was developed by J.B. and U.S. The physical model was built by V.S. with guidance from J.P, A.S. and B.B. The paper was written by J.P. and B.B. with input from all authors.

## Data and materials availability

All data are available in the main text or the supplementary materials. Code used for analysis of the mobile phase in live cells and all the code associated with the model will be made freely available on GitHub. Requests for strains and reagents can be addressed to.

## Extended data

**Figure S1:**
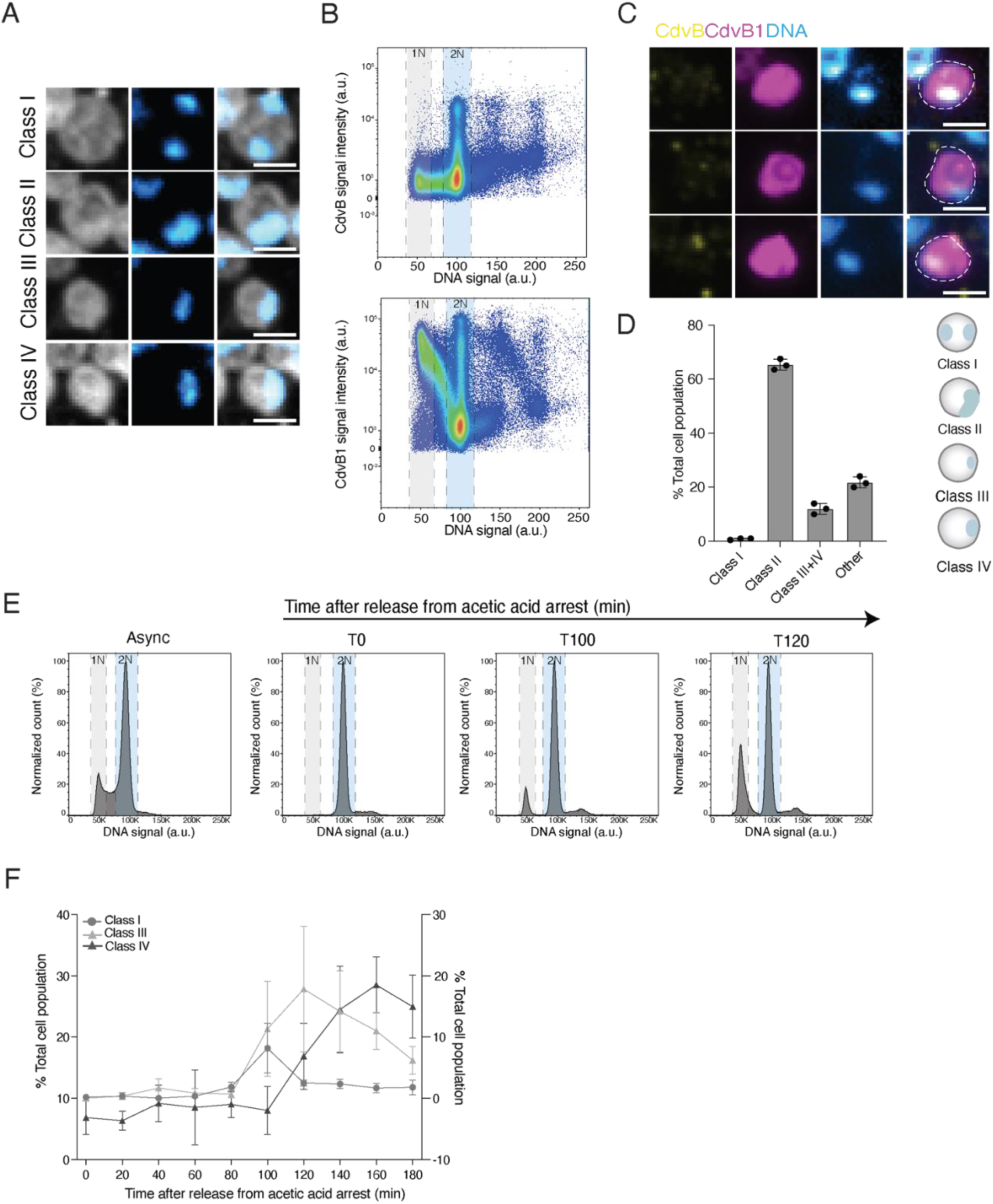
DNA localisation is cell cycle regulated in *Sulfolobus*. **(A)** Panel shows representative images from spinning disc microscopy of wildtype *Sulfolobus* cells labelled with ConA to mark the cell outline (white) and DAPI to label DNA (blue) each of which is used to define a class of cells based on their DNA localisation. Scale bar = 1μm. **(B)** Representative flow cytograms of wildtype cells from asynchronous cultures stained with CdvB and CdvB1 antibodies. Note that CdvB is absent from G1 cells. **(C)** Representative images of immunolabelled G1 cells that have low CdvB and high CdvB1 together with a single, small compact DNA nucleoid. Scale bar = 1μm. **(D)** Quantification of the average proportions of major classes of DNA organisation in fixed cells from asynchronous cultures imaged by spinning disc microscopy defined based on the schematic to the right (n=600, N=3). “Other” represents cells with odd patterns of DNA organization that did not fit into the main categories. **(E)** Representative flow cytograms showing the DNA content of cells at different time points post-release from G2 arrest. **(F)** Proportion of cells with different DNA classes post-release from G2 arrest (n=600, N=3). Class I is plotted on the right Y axis and Classes III and IV are plotted on the left Y axis.

**Figure S2:**
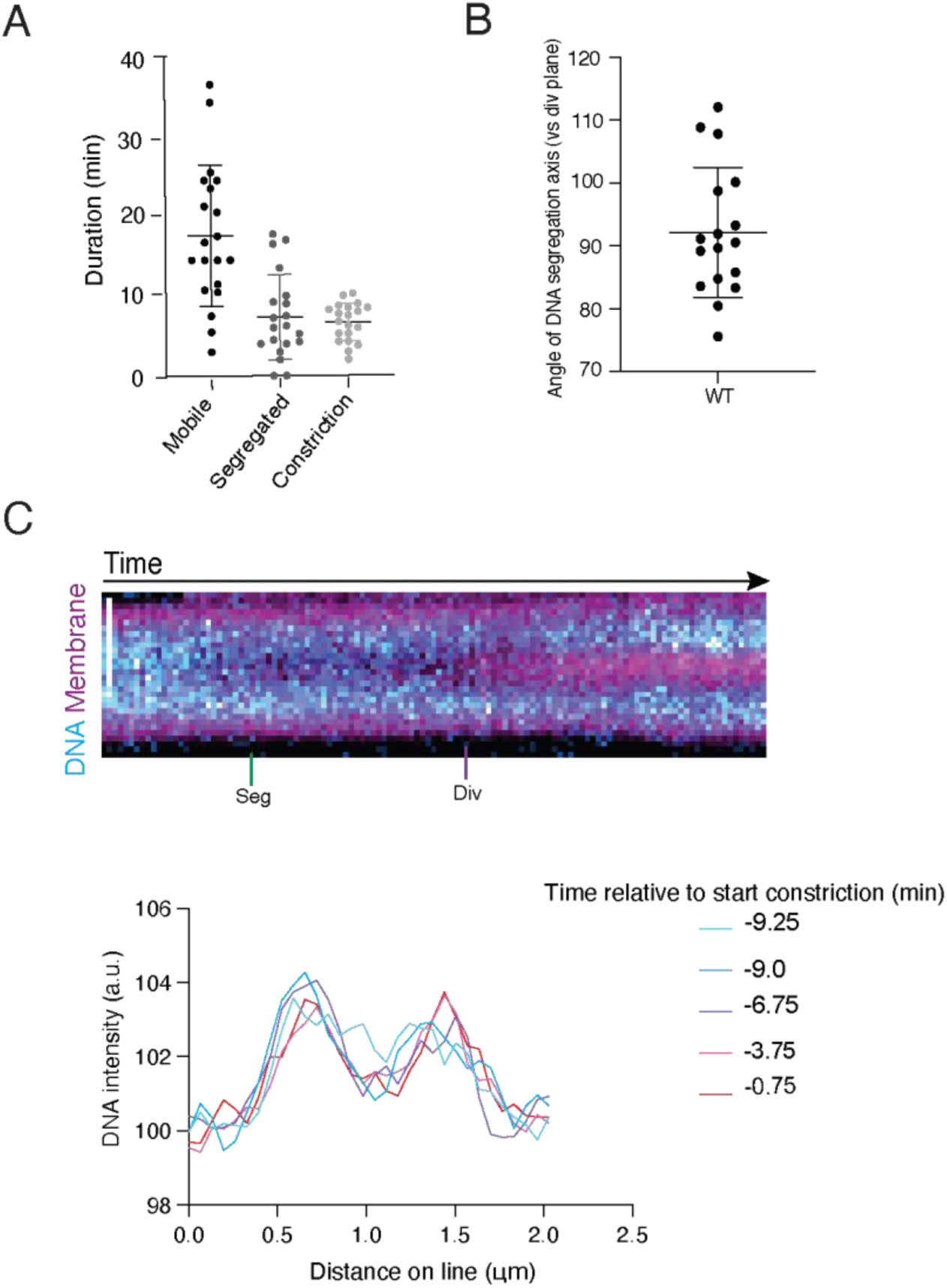
Live imaging reveals DNA dynamics during cell division. **(A)** Plot shows the duration of distinct phases of division (defined in Figure 1) in cells imaged live as they progress from G2 into division. These are defined as: the duration of the mobile phase; the time between DNA segregation and the onset of constriction; and the time between the onset of constriction and its end. (n=20). **(B)** Plot shows a quantification of the angle of DNA segregation axis relative to the plane of cell division (n=17). **(C)** Kymograph shows the DNA and membrane signal across a representative cell imaged live every 15 seconds as it enters division. The DNA segregation event marked with an arrow is rapid event (see arrow) and, after this has occurred, the two spatially separated masses of DNA do not move further apart. Scale bar = 1μm. The corresponding line graph for the kymograph depicted in Fig S2C, showing the DNA intensity profile through a cell from the frame before DNA segregation (t = -9.25 minutes) to just before constriction (t = -0.75).

**Figure S3:**
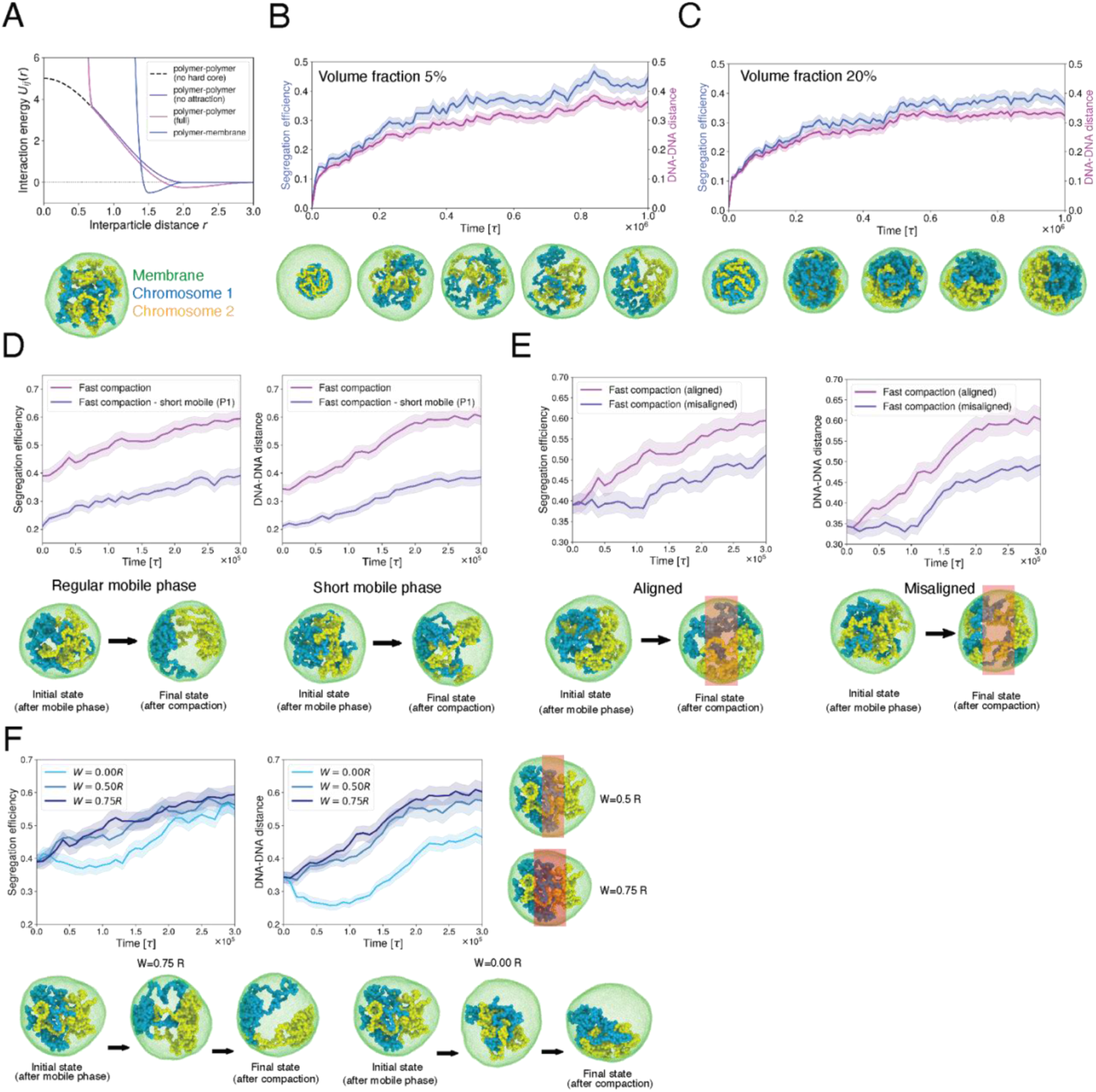
Extended simulation data. **(A)** Interaction potentials between polymer bead pairs and polymer-membrane pairs (see Eq 1-2). The potentials are shown *ɛ*_pp_ ^attr^= 0.25*k_B_T* (full) and *ɛ_pp_* ^attr^= 0 (no attraction) for the polymer-polymer interaction, and *ɛ_pm_*^attr^= 0.5*k_B_T* for the polymer-membrane core. **(B-C)** Segregation efficiency and DNA-DNA distance in simulations of cells undergoing the mobile phase, for the chromosome volume fraction ϕ_p_ ≈ 5%. **(B)** and ϕ_p_ ≈ 20%. **(C).** The snapshots, which represent one of the simulated systems for each ϕ_p_ value, were taken at time intervals of 2.5×10^5^τ. **(D)** Graphs show segregation efficiency (left) and DNA-DNA distance (right) as measured in 50 simulations of cells undergoing fast DNA compaction. Comparison of compaction after a regular-duration mobile phase (duration 10^6^τ) and after a short mobile phase (duration 10^5^τ). The snapshots represent the initial state of compaction, together with compaction at the end of the mobile phase and in the final state, for two of the simulated systems after a regular (left) or short (right) mobile phase. **(E)** Graphs show segregation efficiency (left) and DNA-DNA distance (right) in 50 simulations of cells undergoing fast compaction. Here, we compare the case in which the zone in which compaction is inhibited is perpendicular to the axis of maximum segregation (aligned case) to the one in which there is no alignment (misaligned case). The snapshots represent the initial state of compaction at the end of the mobile phase and the final state reached, for one representative example of the simulated systems for both the aligned and misaligned cases. The red-shaded region defines the area in which compaction has been inhibited. **(F)** Graphs show segregation efficiency and DNA-DNA distance in 50 simulations of cells undergoing fast compaction for different widths *W* of the region in which compaction is inhibited, schematically represented on the right by the red-shaded regions. The snapshots, taken at intervals of 1.5×10^5^τ, show two simulated systems undergoing compaction for *W*= 0.75*R* and *W*= 0. The latter value corresponds to uniform compaction. For graphs Fig S3B-F, n=50, and the shaded area represents the standard error. In all the snapshots, the cell membrane is coloured in green, while the two chromosomes are coloured in yellow and blue, respectively.

**Figure S4:**
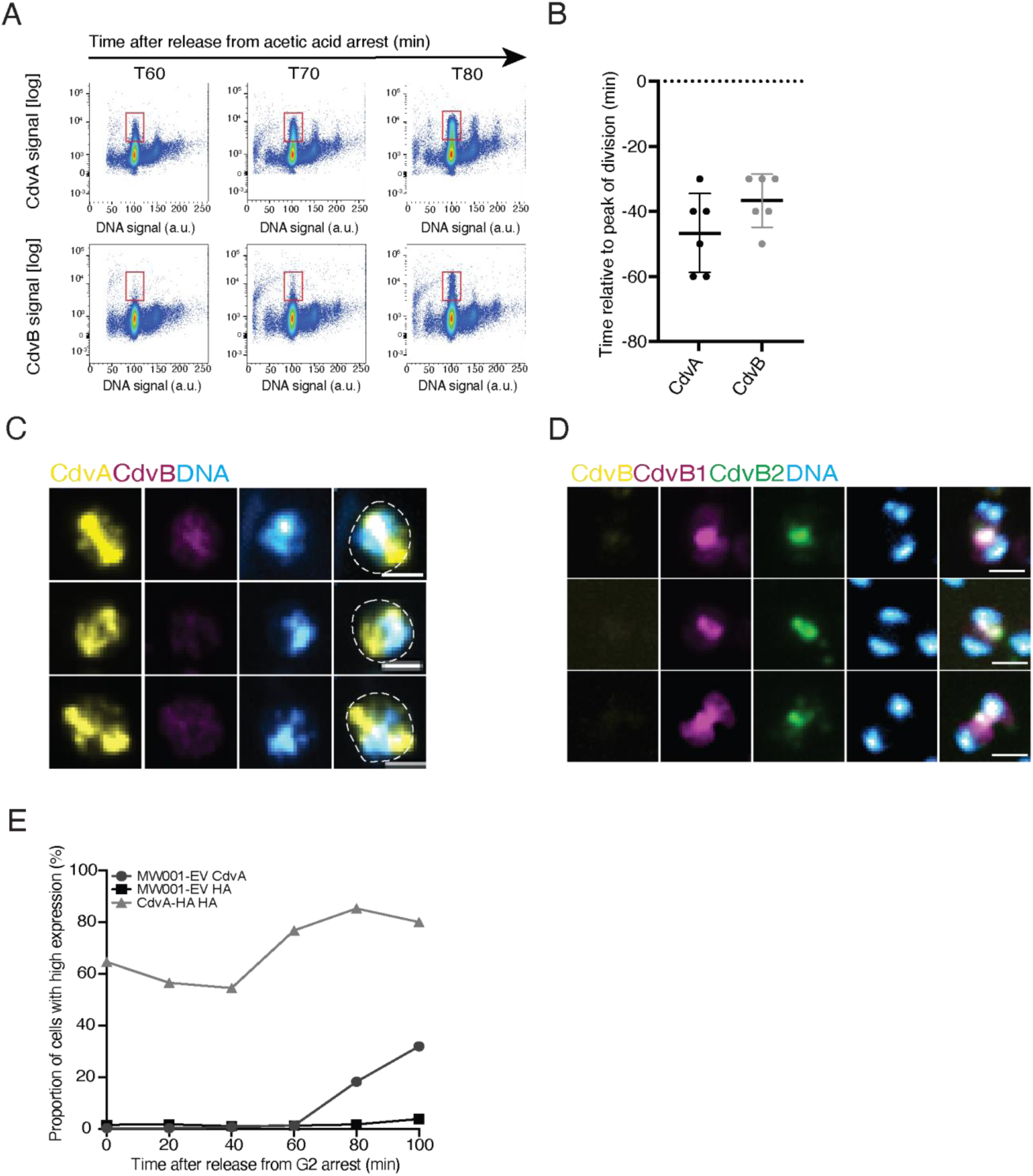
Ring states through cell division phase. **(A)** Representative flow cytograms of control (MW001) cells at sequential time points post-release from a G2 acetic acid arrest that have been fixed and stained for CdvA and CdvB. **(B)** Plot shows first observable expression of CdvA and CdvB following the release of wildtype (DSM639) cells from a G2 arrest as quantified from cytograms relative to the peak of division (t=0). N=6 **(C)** Representative images showing DNA in DSM639 cells with ring-like CdvA structures before the accumulation of ESCRT-III polymers. **(D)** Representative images of DNA in DSM639 cells in late division as measured by the presence of contracting CdvB1 and CdvB2 rings following the removal of CdvB from the ring. **(E)** Quantification of CdvA-HA induction in an early induced overexpression during an acetic acid arrest. Scale bars =1μm.

**Figure S5:**
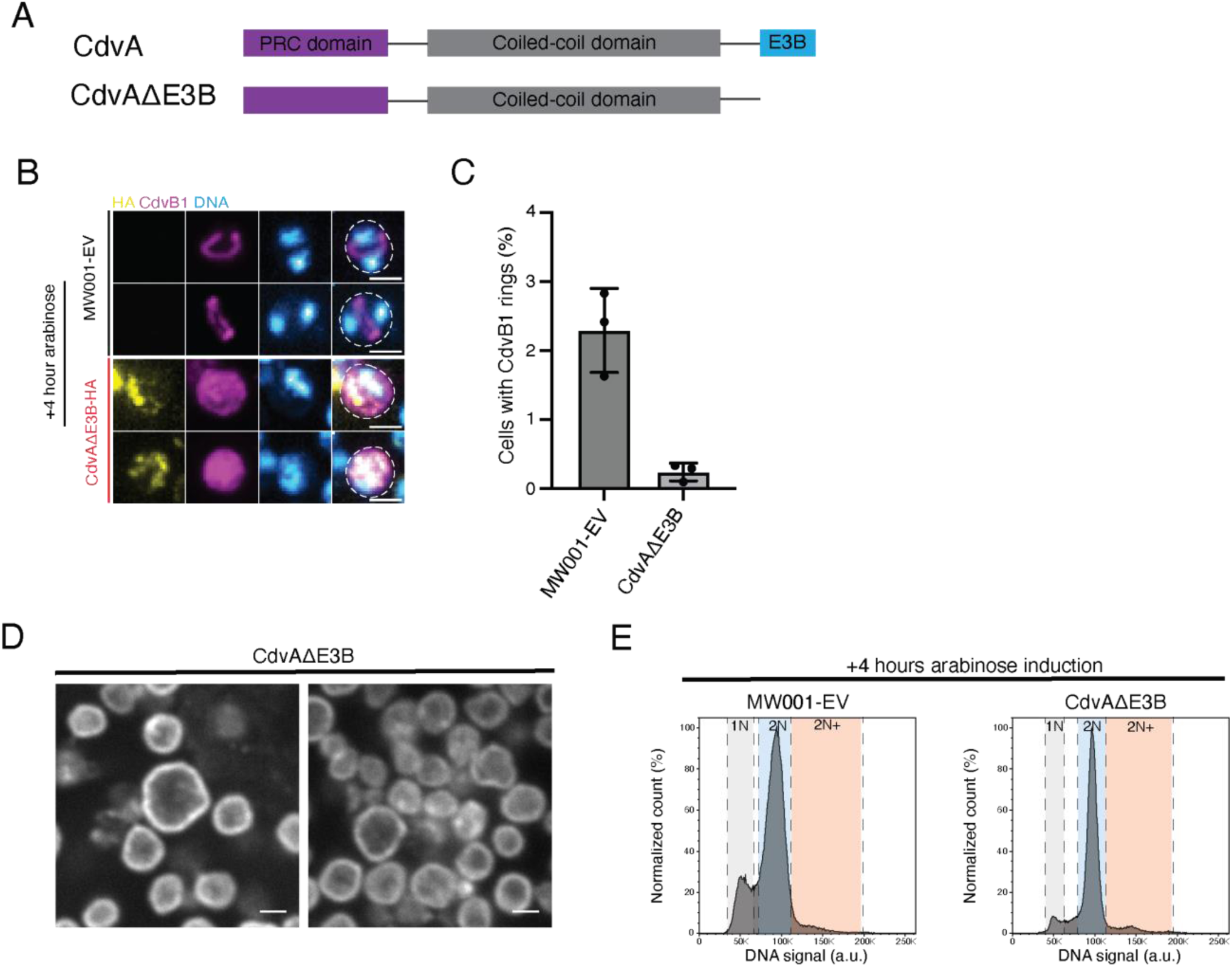
CdvAΔE3B blocks ESCRT-III ring formation and cell division. **(A)** Schematic of the full length CdvA protein and the CdvAΔE3B truncation mutant. **(B)** Representative immunofluorescent images of MW001-EV and CdvA^ΔE3B^-HA cells stained with HA and CdvB1 after 4 hours of arabinose induction. Scale bar = 1μm. **(C)** Quantification of proportion of cells that have CdvB1 rings in MW001-EV and CdvAΔE3B-HA after 4 hours arabinose induction (n=>2500, N=3). **(D)** Immunofluorescent images of large CdvA^ΔE3B^-HA cells stained with ConA that have failed multiple rounds of cell division. Scale bar 1μm. **(E)** Flow cytograms of MW001 cells carrying an empty vector and those carrying a plasmid encoding CdvAΔE3B-HA after 4 hours of arabinose induction, showing a clear reduction in the G1 population in cells expressing CdvAΔE3B.

**Figure S6:**
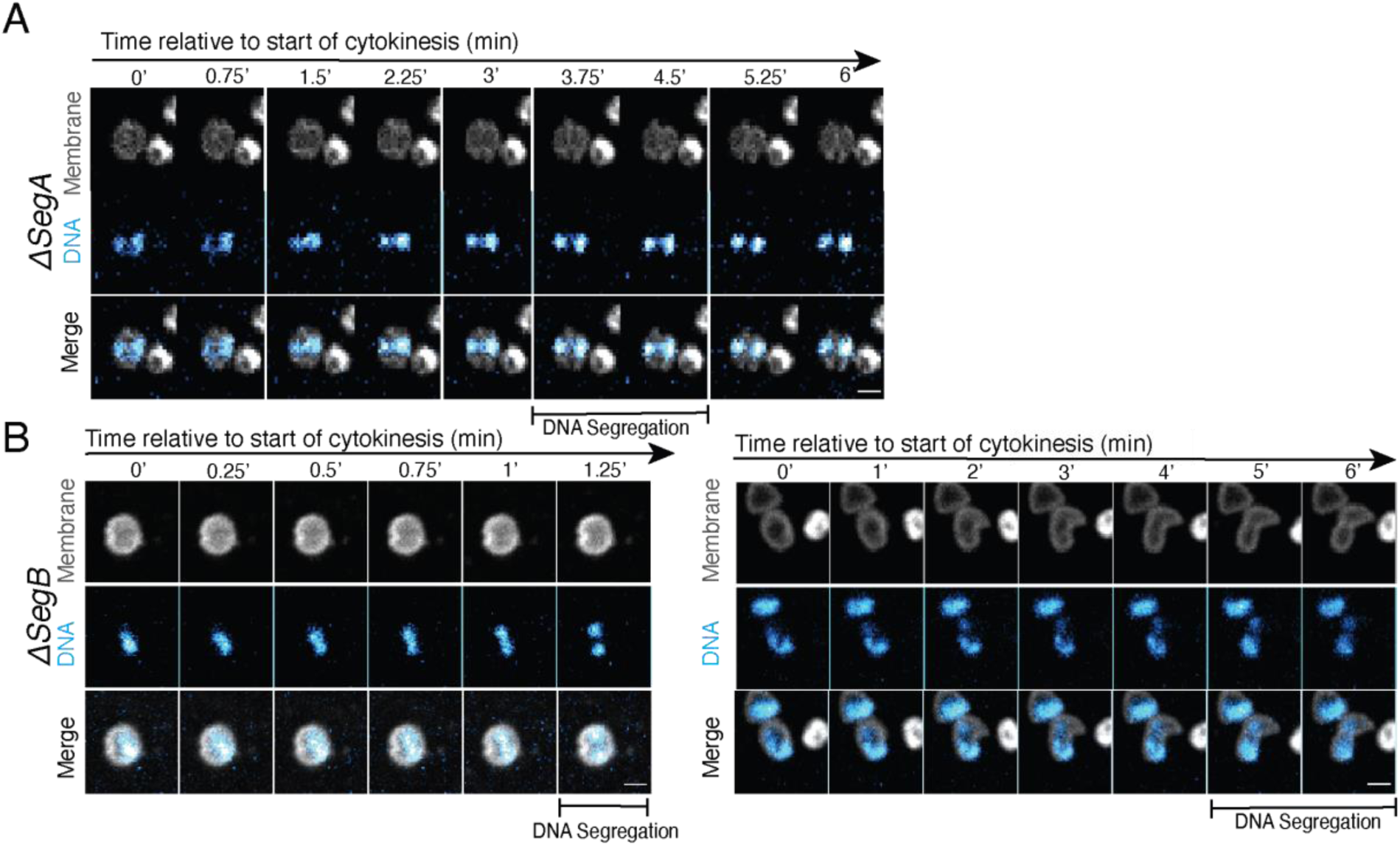
SegA and SegB have a roll in DNA compaction. **(A)** Representative montage from live imaging of Δ*SegA* cells with late DNA segregation. **(B)** Representative montage from live imaging of Δ*SegB* cells with late DNA segregation and no compaction error (left) and late DNA segregation with significant DNA compaction error (right). Scale bars = 1μm.

**Figure S7:**
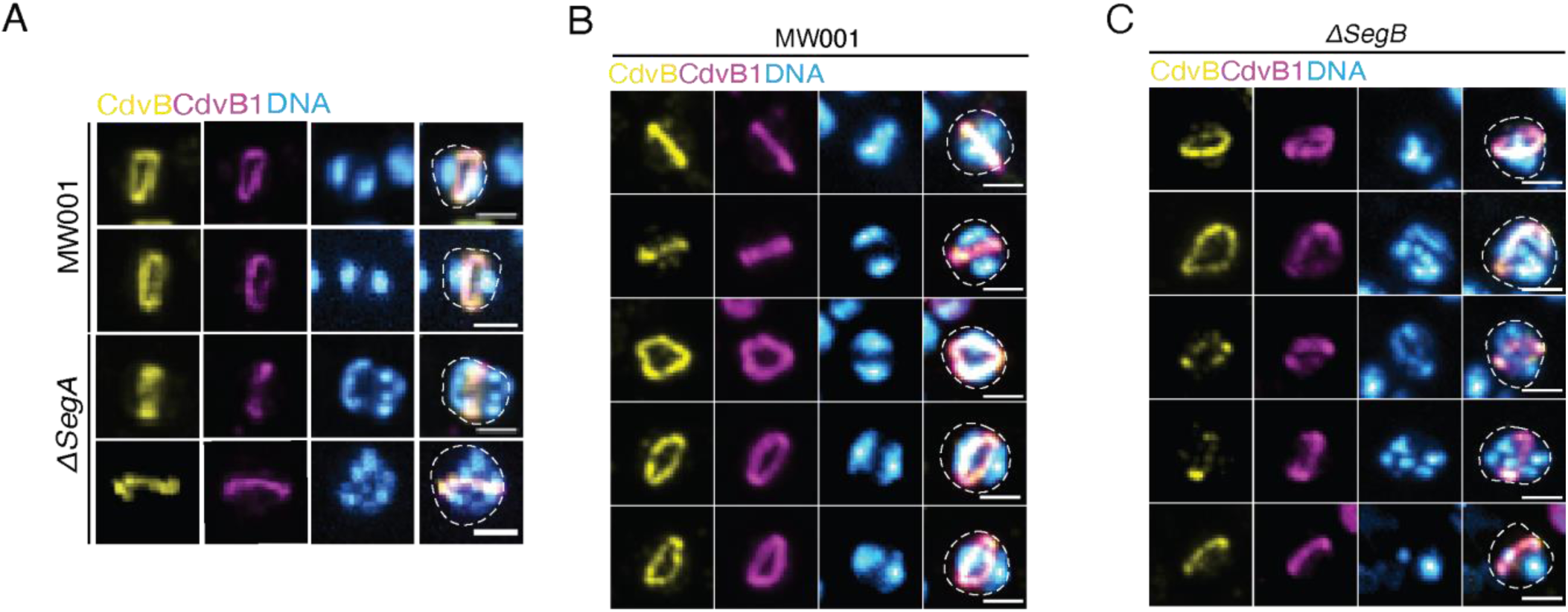
SegA and SegB facilitate DNA compaction at division. **(A)** Representative immunofluorescence images of MW001 and Δ*SegA* cells with CdvB+CdvB1 rings. **(B)** Representative immunofluorescence images of MW001 cells with CdvB+CdvB1 rings. **(C)** Representative immunofluorescence images of Δ*SegB* cells with CdvB+CdvB1 rings. Scale bars = 1μm.

**Figure S8:**
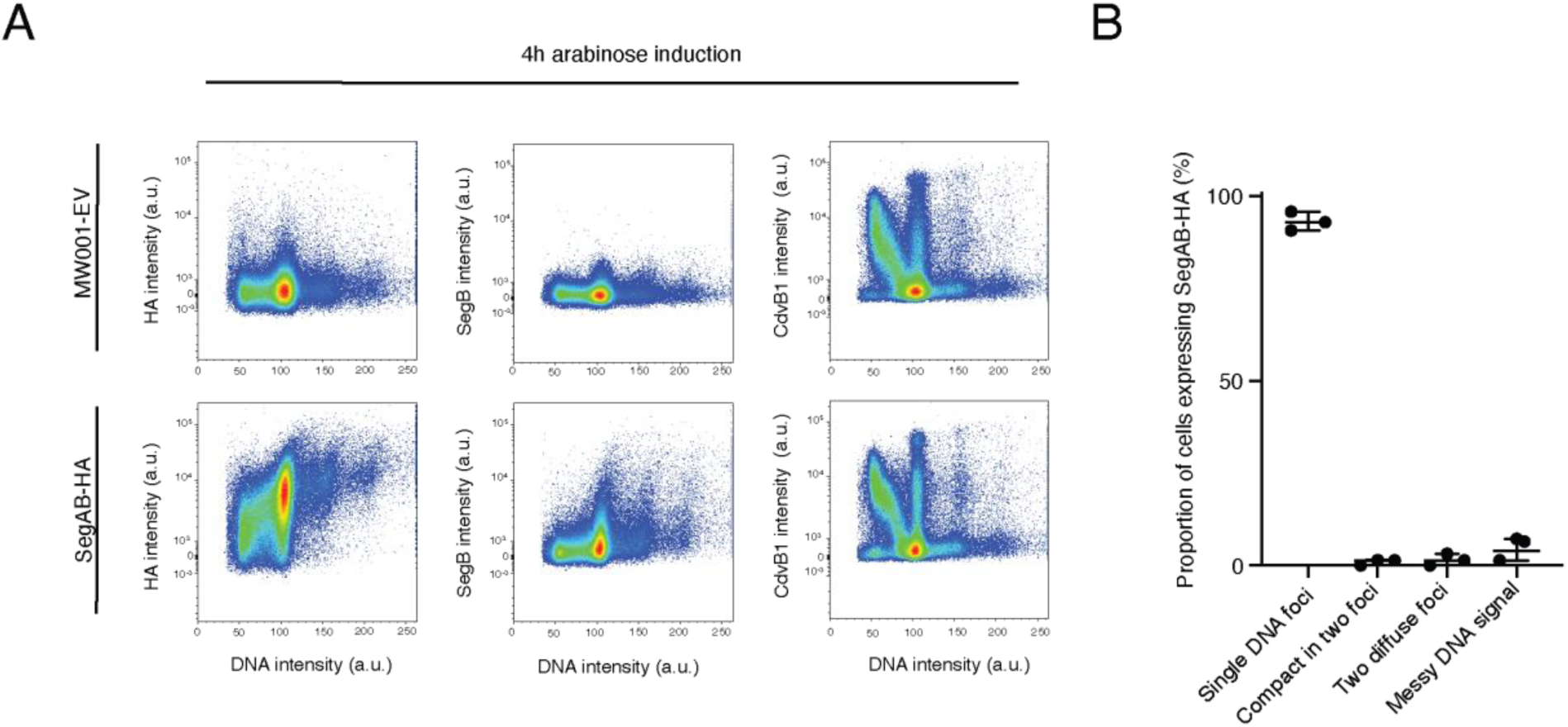
SegAB overexpression isn’t sufficient to induce DNA segregation. **(A)** Flow cytograms showing the HA signal in MW001-empty vector and SegAB-HA expressing cells after 4 hours of treatment with arabinose. **(B)** Quantification of DNA segregation in cells that lack CdvB1 but which express SegAB-HA (n=337, N=3)

## Supplementary tables

**Table S1:** List of primary antibodies.

| Antibody (animal) | Supplier |
| --- | --- |
| Anti-CdvA (rabbit) | Lab generated |
| Anti-CdvA (chicken) | Lab generated |
| Anti-CdvB (rabbit) | Lab generated |
| Anti-CdvB (rat) | Lab generated |
| Anti-CdvB1 (chicken) | Lab generated |
| Anti-CdvB2 (guinea pig) | Lab generated |
| Anti-SegB (rabbit) | Dyche Mullins lab |
| Anti-HA (mouse) | ThermoFisher, 26183 |

**Table S2:** List of secondary antibodies.

| ThermoFisher | Anti-Rabbit | Anti-Chicken | Anti-Guinea pig | Anti-Mouse | Anti-Rat |
| --- | --- | --- | --- | --- | --- |
| Alexa Fluor 488 | A11034 | A11039 | A11073 | A10631 | A11006 |
| Alexa Fluor 546 | A11035 | A11040 | A11074 | A11030 | A11081 |
| Alexa Fluor 647 | A21245 | A21449 | A21450 | A21235 | A21287 |

## Materials and Methods

### Strains, culture media and growth conditions

*S.acidocaldarius* strains DSM639 (WT) MW001-CdvA (MW001 with the full length and ΔE3B domain truncation), MW001-SegAB (MW001 with the SegAB overexpression plasmid) and MW001-EV (MW001 with the empty vector plasmid) were grown at 75°C, pH 3.0-3.5, in Brock medium supplemented with 0.1% w/v NZ-amine and 0.2% w/v sucrose. The mutant *S.acidocaldarius* strains MW001 (uracil auxotroph, genetic background), MW001-Δ*segA* (MW001 with *segA* gene deletion), MW001-ΔSegB (MW001 with SegB gene deletion), MW001-ΔSegAB (MW001 with SegAB gene deletion) and were supplemented with 4μg/ml uracil. The optical density of liquid cell cultures was maintained at levels corresponding to exponential growth, OD600 values between 0.100 and 0.400, for between 2-4 days prior to experiments and samples were only taken within this range. Cells were fixed by stepwise addition of absolute ethanol until a concentration of 70%.

### Cell Synchronization

*S.acidocaldarius* cultures were arrested by treatment with acetic acid (final concentration 3mM) for 4 hours. After arrest, cells were washed 3 times in fresh Brock medium to remove the acetic acid and released into fresh media to re-enter the cell cycle. Cells were then fixed as previously described at different time points post-release.

### Molecular genetics

Deletion mutants were constructed using the pSVA406 vector as described previously^1^. Briefly, the upstream and downstream regions of a gene of interest was amplified via PCR and cloned into pSVA406 through restriction digest. Positive clones were selected through analytical digestion, methylated and purified via miniprep for transformation into electrocompetent *S. acidocaldarius* cells. Positive strains containing the integrated inserts were next re-streaked onto Gelrite-Brock plates supplemented with 4μg/mL uracil and 100μg/mL 5-fluoroorotic acid (5-FOA; Zymo Research, P9001-1). Colonies obtained were then selected, grown, and screened for the deletion of the gene of interest through genomic DNA extraction and genotyping of the overlapping regions of the locus of interest. Positive clones were frozen in Brock medium containing 50% glycerol (v/v) and stored at -70°C.

The SegAB deletion mutant was constructed by the deletion of the entire operon from the start codon of SegA to the stop codon of SegB inclusive. Due to the 4 nucleotide overlap between the SegA and SegB genes, the single Seg mutants were constructed by deletion of the gene of interest without affecting the neighbouring gene: the SegA deletion mutant was constructed by the deletion of the gene from the start codon to before the start codon of SegB; the SegB deletion mutant was constructed by the deletion of the gene from after the stop codon of SegA to the stop codon of SegB.

To generate the overexpression plasmid of SegAB with C-terminal HA tag, Saci_0203+Saci_0204 (uniport accession number Q4JC57 and Q4JC56) was PCR amplified from the genomic DNA of *S. acidocaldarius* DSM639 using the primer pair: 5’-AATccatggctATGATAATCACTGTCATCAA-3’and 3’CGCctcgagACTCTTTTTACTCTCTAATG-5’ and cloned into the plasmid vector containing arabinose-inducible promoter (pSVAaraFX-HA). The PCR product was purified by Monarch DNA cleanup kit (New England Biolabs) and the vector was digested with restriction enzymes NcoI and XhoI followed by gel extraction. The linearized vector and purified PCR product were assembled by Gibson assembly using the overlapping sequence. The sequence of the cloned plasmid (pSVAaraFX-SegAB-HA) was verified by Sanger sequencing.

To generate the overexpression plasmid of CdvAΔE3B with C-terminal HA tag, 1 to 220 aa of saci_1374 (uniport accession number Q4J923) was PCR amplified from the genomic DNA of *S. acidocaldarius* DSM639 using the primer pair: 5’-taataattgataagcgtcttacttatcataccATGGGCATTCCGGTTGA-3’ and 5’-tacgcgtagtccggaacgtcatacgggtactcgagCTCGTTCTTATTTGACTGTTCTGTTG-3’ and cloned into the plasmid vector containing arabinose-inducible promoter (pSVAaraFX-HA). The PCR product was purified by Monarch DNA cleanup kit (New England Biolabs) and the vector was digested with restriction enzymes NcoI and XhoI followed by gel extraction. The linearized vector and purified PCR product were assembled by Gibson assembly using the overlapping sequence flanking the CdvAΔE3B (lower case region of the primers above). The sequence of the cloned plasmid (pSVAaraFX-CdvAΔE3B-HA) was verified by Sanger sequencing.

### Live cell imaging of *S. acidocaldarius*

Live cell imaging was performed using the Sulfoscope set-up as described^2^ with additional hardware modifications detailed^3^. Briefly, Attofluor chambers (Invitrogen A7816) were assembled with 25mm coverslips and filled with 300µl BNS media. The media was allowed to dry onto the surface of the coverslip at 75° C before the chambers were washed thoroughly with BNS and placed into the pre-heated Sulfoscope and allowed to equilibrate to 75° C. 5ml of *S. acidocaldarius* cell culture at an optical density at 600 nm (OD600nm) of 0.15 to 0.3 was kept at 75° C and stained with CellMask Deep Red Plasma Membrane (Invitrogen C10046; 1:5000). 400µl of the stained cell suspension was transferred to the pre-heated Attofluor chamber, making sure to avoid cooling of the chamber or the cells. Cells were immobilised using semi-solid Gelrite pads (0.6% Gelrite, 0.5× BNS pH 5, 20 mM CaCl2). Gelrite pads were prepared by mixing thoroughly pre-heated BNS (pH 5) and 1.2% Gelrite in a 1:1 ratio. To set, 15ml of the molten Gelrite BNS solution was added to 9cm plastic petri dishes and allowed to set at room temperature (∼5 minutes). To prepare the immobilsation pads, half-moon shapes were cut from the plate with a 7mm diameter circle punch and placed onto 13-mm circular coverslips. Immediately before imaging pads were incubated at 75°C for 5min in a bead bath until they equilibrated to the imaging temperature as well as began to dry causing the edges of the pad to curve downwards. Upon adding the cell suspension, the pre-heated immobilisation pad was placed such that the concave edge of the pad was in the center of the chamber. Cells were imaged at this concave edge where diffusion is limited but cells are not subjected to mechanical stress by the pad. Images were acquired on a Nikon Eclipse Ti2 inverted microscope equipped with a Yokogawa SoRa scanner unit with an additional x2.8x magnification and Prime 95B sCMOS camera (Photometrics). Imaging was performed with a 60× oil immersion objective (Plan Apo 60×/1.45, Nikon) using a custom formulated immersion oil for high temperature imaging (maximum refractive index matching at 70°C, n = 1.515 ± 0.0005; Cargille Laboratories). Images were acquired at a t1010otal magnification of ×168 with 15-ms exposure time and 10% laser power at intervals of 15 seconds for 2.5 hours or until any cell death was observed. XY drift was corrected after acquisition using the ImageJ plugin StackReg^4^.

### Live imaging quantification

As a first step, time-lapse microscopy image sequences were manually cropped using Fiji to isolate regions of interest containing individual cells. Each cropped sequence was subsequently processed via a custom Python script to segment cellular and DNA components, quantify apparent motion, and visualize the results. Prior to segmentation, images underwent pre-processing, consisting of a spatiotemporal Gaussian filter and deblurring via ten iterations of the Richardson-Lucy algorithm^5,6^. Cellular segmentation and tracking were performed using the “cyto2” model implemented in Cellpose^7^, employing a diameter parameter of 22 and utilizing both available channels. In instances where the resulting masks were null, a watershed segmentation approach was employed as an alternative. The centroid of the resulting mask was then tracked using nearest-neighbour assignment, with other segmented regions being discarded. Apparent motion, or optical flow, was calculated for each channel using the Lucas-Kanade algorithm^8^ as implemented in scikit-image^9^. DNA signal segmentation was achieved using a difference of Gaussians filter, followed by temporal tracking using the Crocker and Grier algorithm^10^ as implemented in trackpy^11^. Processed images, segmentation masks, and calculated flow fields were then stored in HDF5 format. Subsequently, frame difference, momentum, and momentum divergence were computed and visualized as streamlines. Furthermore, the mean of the magnitude of each of these three derived quantities, calculated within the segmented cellular regions, was determined at each time point to quantitatively assess DNA motion over time.

### Immunolabelling

Fixed cells were washed and rehydrated in PBS-TA (PBS supplemented with 0.2% Tween20 and 3% bovine serum albumin) before incubation overnight at 25°C and 400 revolutions per minute (rpm) agitation with primary antibodies (table S1). Conjugated secondary antibodies were used for detection of target proteins (table S2) by staining for 2-3 hours at 25°C and 400 rpm agitation. The S-layer was stained by incubating the rehydrated sample with 200 μg/ml Concanavalin A conjugated to Alexa Fluor 647 (ThermoFisher, C21421) for 2-3 hours. DNA was visualized by the addition of 1 μg/ml DAPI (4′,6-diamidino-2-phenylindole; Thermo Fisher Scientific, 62248) to samples after secondary antibody incubation. For spinning disc microscopy, Lab-Tek chambered slides (Thermo Fisher Scientific, 177437PK) were coated with 2% polyethyleneimine (PEI) for 30 minutes at 37°C. Chambers were washed with Milli-Q water before stained cell suspension was added and spun down for 1 hour at 750 relative centrifugal force (RCF).

### Spinning disc microscopy

Cells were imaged in Lab-Tek chambered coverslip using a Nikon Eclipse Ti2 inverted microscope equipped with a Yokogawa SoRa scanner unit and Prime 95B scientific complementary metal-oxide semiconductor (sCMOS) camera (Photometrics). Images were acquired with a 100× oil immersion objective (Apo TIRF 100×/1.49, Nikon) using immersion oil (immersion oil type F2, Nikon). A total magnification of ×280 was achieved using the ×2.8 magnification lens in the SoRa unit. Images were acquired with 200-ms exposure time for labelled proteins and 500-ms exposure time for DNA stains, with laser power set to 20%. *z*-axis data were acquired using 10 captures with a 0.18-0.22-μm step.

### Flow cytometry

DNA was labelled with 2μM Hoescht for flow cytometry. Cells were gated by DNA staining (UV excitation). Laser excitation wavelengths of 355, 488, 561 and 633nm were used in conjunction with the emission filters 450/50, 530/30, 586/15 and 670/14 respectively. Flow cytometry analysis was performed on BD Biosciences LSRFortessa. Side scatter and forward scatter was recorded. Analysis was performed using FlowJo v10.8.1.

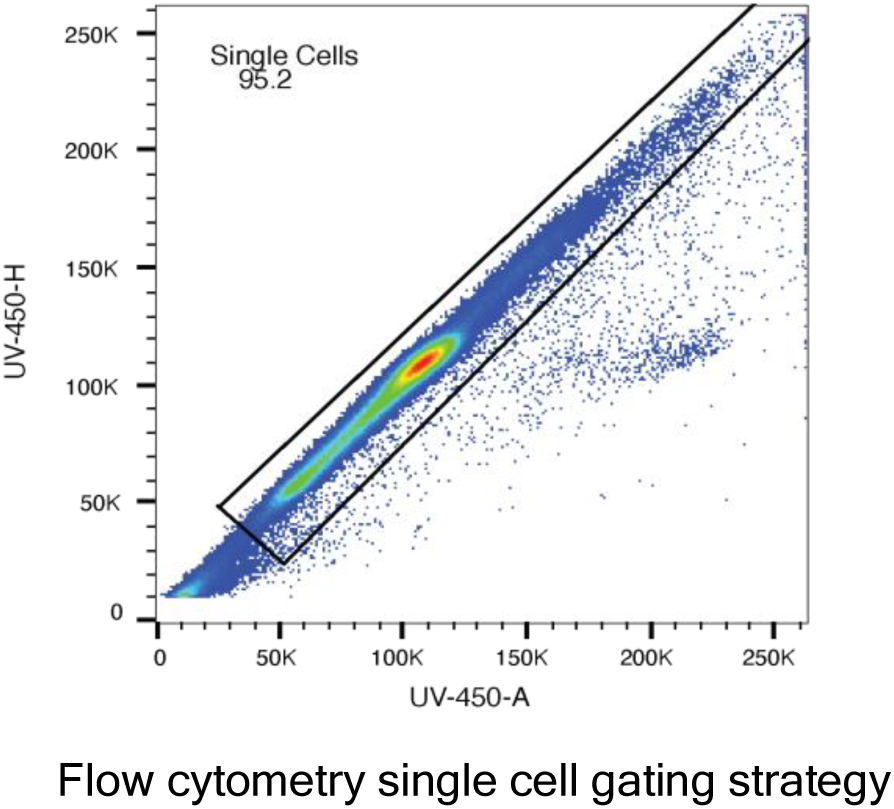

### Statistical analysis

All statistical analysis was performed in Microsoft Excel or GraphPad Prism 10 software. Significance was defined as p ≤0.05. Significance levels used were *p ≤0.05, ** p ≤0.01, *** p ≤0.001 and **** p ≤0.0001. Exact statistical tests are reported in the figure legends.

## Detailed description of the computational model

### Modelling interaction potentials

The two chromosome copies were modelled as bead-spring ring polymers of *N*chr = 500 beads each. Since the genome of *Sulfolobus* comprises *≈* 3 Mbp^12^, each bead represents a chromosome section of size *≈* 6 Kbp. These values are in the same range as those commonly used in coarse-grained models of bacterial chromosomes^13–18^. Polymer segments are allowed to cross with a small energy penalty to simulate the action of topoisomerase-II-like enzymes^19–22^. Additionally, the two chromosomes are enclosed in a thin fluid vesicle modelling the plasma membrane^23^. The model used for the vesicle beads follows Yuan *et al*^23^, which consists of a one-bead-thick, solvent-free fluid membrane, we use *r*min= 2^1*/*6^*σ*, *rc*= 2.6*σ*, *ɛ*= 4.3*kBT*, *ζ*= 4, *µ*= 3, *θ*= 0, *β*= 0. This choice of parameters ensures that the membrane is fluid. Here, *σ* is the diameter of a membrane bead, which we take as the unit of length. Throughout this work, we additionally take as units of energy and mass respectively *kBT* (with *kB* Boltzmann’s constant and *T* the absolute temperature) and *m*, the mass of a membrane bead.

Each pair of polymer beads interact with a potential

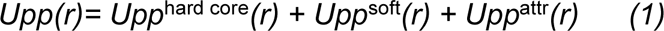

where: *U_pp_*^hard core^(*r*) is a hard-core repulsion term^24^ introduced to prevent a complete collapse of the polymer chain in the presence of attractive interactions; *U*_pp_ ^soft^(*r*) is a soft repulsive potential, which allows chain crossing^19–22^; and *U_pp_* ^attr^(*r*) is a short-ranged attractive interaction^25^. Here *r* is the interparticle distance. Similarly, polymer and membrane beads interact with the potential

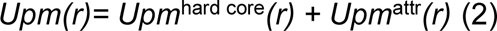

These potentials have the following general expressions for two pair of beads *i, j*, where *i, j*= *p,m* for polymer and membrane beads, respectively:

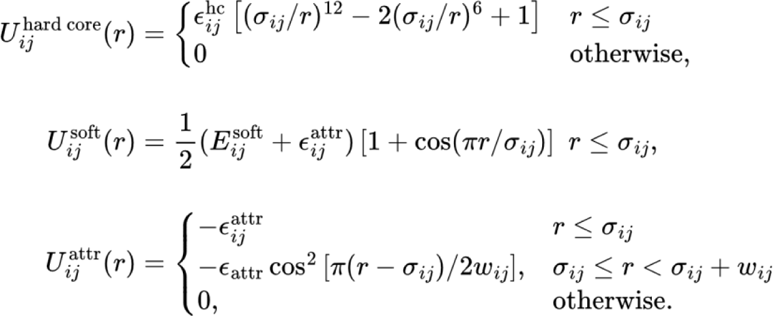

For the polymer-polymer (*pp*) interactions, the parameter values are *σ_pp_*= 2*σ*, *ɛ*_pp_ ^hc^= 4*k_B_T*, *E*_pp_ ^soft^= 5*k_B_T*, and *w_pp_*= *σ* (width of attractive potential well). The parameter *ɛ*_pp_ ^attr^ tunes the polymer-polymer attraction, and thus the degree of compaction of each chromosome. We note that the attraction is between any pair of beads, irrespective of which chromosome they belong to. The mass of a polymer bead is *m_p_*= 1.1*m*, slightly larger than the mass of a membrane bead.

For the polymer-membrane (*pm*) interactions, the parameter values are *σ_pm_*= (*σ_pp_* + *σ*)*/*2= 1.5*σ*, *ɛ_pm_*^hc^= 4*k_B_T*, and *w_pm_*= 0.5*σ*. The parameter *ɛ_pm_*^attr^ tunes the attraction between polymer and membrane beads, resulting in association of the polymers to the membrane; in what follows, we set *ɛ_pm_*^attr^= 0.5*k_B_T*. The polymer-polymer and polymer-membrane potentials are shown in (Fig S3A) for the values *ɛ*_pp_ ^attr^= 0.25*k_B_T* and *ɛ_pm_*^attr^= 0.5*k_B_T*. Bonded neighbours in the same polymer chain additionally interact with an harmonic potential, *U*_bond_(*r*)= *K*(*r − r*_0_)^2^, where *K*= 100*k_B_Tσ^−^*^2^ is the bond strength and *r*_0_= 0.7*σ_pp_=* 1.4*σ* its equilibrium length.

### Simulations

The initial state of the simulation is obtained by taking the two ring polymers (chromosomes), with each bead in a ring connected to the corresponding one in the other ring by a harmonic bond (so that the two polymers are in a “ladder-like configuration), and compressing them into a sphere until they reach volume fraction 40%. Then, the bonds between the two different rings are removed, while the intra-ring bonds are left untouched. This protocol ensures that there is initially no topological linking between the two rings. However, if linking was initially present, it would quickly be lost, since we allow for (intra- and inter-) chain crossing. We generate *n*_repeat_= 50 initial configurations, which result in 50 simulation repeats for every set of parameters. The two ring polymers are then enclosed in a spherical vesicle. For most of this work, the number of vesicle beads is *N*mem= 6750, resulting in a mean radius *R*= 21.4*σ* and a chromosome volume fraction *ϕ_p_ ≈* 10% inside the vesicle (Fig S3A). However, we also consider vesicles of *N*mem= 10000 beads (*R*= 26.0*σ*, *ϕ_p_ ≈* 5%) and *N*mem= 4250 beads (*R*= 17.1*σ*, *ϕ_p_ ≈* 20%).

Following the creation of the initial configuration, each simulation was broken up into two discrete phases, P1 and P2: P1 corresponds to the experimentally observed “mobile phase”, while P2 corresponds to chromosome compaction. Below, we briefly describe these phases. P1 (mobile phase): During this phase, the system is simulated with no polymer-polymer attraction (*ɛ*_pp_ ^attr^= 0) and no polymer-membrane association (*ɛ_pm_*^attr^= 0). We let the system evolve for a time *T*_1_= 10^6^*τ*, with *τ*= (*mσ^2^/k_B_T)^1/2^* the simulation unit of time, unless otherwise specified. Since during P1 there are no attractive interactions between the chromosomes, the only driving force for segregation is the entropic penalty resulting from the overlap of polymeric coils^13–15,26–28^ As discussed in the main text, this force is sufficient to lead to an appreciable degree of segregation. P2 (compaction): During P2 we introduce both compaction and membrane association by setting the polymer-polymer attraction to *ɛ*_pp_ ^attr^= 0.25*k_B_T* and the polymer-membrane one to *ɛ_pm_*^attr^= 0.5*k_B_T*.

The total duration of P2 is 30% that of P1, *i.e.*, *T*_2_= 3*×*10^5^*τ* . Whereas membrane association is introduced gradually in all cases, by increasing *ɛ_pm_*^attr^ over *T*_2_*/*2 time steps, we consider two protocols to introduce compaction. These are fast compaction, in which *ɛ_pp_*^attr^ increases suddenly from 0 to the final value over a very short time *T*_2_*/*30= 10^4^; and slow compaction, in which it increases gradually over the same timescale as the one chosen for membrane association, *i.e. T*_2_*/*2. Since we observe experimentally that compaction happens away from the mid-cell, we only allow compaction outside of the region *L_x_/*2 *− W/*2 *< x < L_x_/*2 + *W/*2, with *L_x_* the length of the simulation box in the *x* direction. Unless otherwise stated, the *x* direction in P2 is chosen to match with the direction of the maximum segregation axis, derived from Linear Discriminant Analysis applied to the final configuration of P1 (see *Quantification of segregation* below). Thus, the region in which compaction is inhibited lays perpendicular to the segregation axis. The parameter *W* represents the width of the “no-compaction” region in which compaction is inhibited: in practice, for all polymer beads in this region *ɛ*_pp_ ^attr^ is set to zero. In this work, we take *W*= 0.75*R*. We also test *W*= 0.5*R*, finding that our results remain largely unaffected for this lower value. For *W=* 0 (uniform compaction), instead, the amount of segregation is significantly lower.

The simulations are performed at constant number of particles, volume and temperature. The solvent is treated implicitly using a Langevin thermostat^29^, which also ensures that the temperature *T* is kept constant. The viscous friction that each bead experiences is *ζ*= 10*m/τ*. The simulations are carried out using LAMMPS^30^, where time integration is performed using the velocity Verlet algorithm, with time step *δt*= 10*^−^*^2^*τ*. Periodic boundary conditions are applied in all three spatial directions, however this is irrelevant except for a few membrane particles very rarely being released from the vesicle.

### Quantification of segregation

To quantify the amount of segregation of the two chromosome copies, we employ two different methods. In the first and most straightforward case, the distance between the centres of mass (CoM) of the two chromosomes is measured (which we henceforth call DNA-DNA distance for simplicity). We normalize this quantity by dividing it by the mean vesicle radius *R*; this also facilitates the comparison between vesicles of different sizes. The second method we employ is Linear Discriminant Analysis (LDA)^31^, which we implement using the Python module scikit-learn^32^. In practice, this amounts to trying to find a 2D plane in 3D space such that all the coordinates of the first chromosome lay to one side of the plane,with all the coordinates of the second chromosome laying to the other side. The normal vector to this plane defines what we call the *segregation axis*. If the two chromosomes are perfectly segregated, finding a plane that exactly separates their coordinates will be possible. In general, however, this procedure will result in a certain fraction *f* of the polymer coordinates being “misclassified”, *i.e.*, laying on the wrong side of the best-fitting plane. Since for perfectly mixed coordinates *f= 1/2* (50% of the coordinates are misclassified), we define a segregation efficiency *s= 1 − 2f*, which is equal to 0 for perfect mixing (*f= 1/2*) and to 1 for perfect segregation *(f= 0)*.

### Effect of chromosome volume fraction on segregation

In Fig 1E we showed the extent of segregation during the mobile phase, quantified both by the segregation efficiency and the DNA-DNA distance between the two chromosomes (see *Quantification of segregation*). The chromosome volume fraction considered in the main text is *ϕ_p_ ≈* 10%, which is the one that best matches the one estimated from visual inspection of the experimental images. However, since it is difficult to give a precise estimate of the volume fraction of the chromosome of *Sulfolobus*, we check here that our results are robust with respect to variations in the chromosome volume fraction. We show the results of simulations performed for *ϕ_p_≈* 5% (Fig S3B) and *ϕ_p_≈* 20% (Fig S3C). We observe that the extent of segregation reached at the end of the mobile phase increases slightly when the volume fraction decreases; however, the overall phenomenology remains the same. We thus conclude that our results are robust with respect to changes in the chromosome.

### Effect of mobile phase duration

In the main text, we set the duration of the mobile phase to be long enough that the system reaches equilibrium, and that the extent of segregation (as measured by the segregation efficiency and the DNA-DNA distance) reaches on average a plateau. We also investigated the impact of shortening the duration of the mobile phase to 10% of its normal duration (10^5^τ instead of 10^6^τ). The results are shown (Fig S3D) where we compare the segregation efficiency and the DNA-DNA distance during fast compaction for the two cases. As expected, the initial extent of segregation is significantly lower after a shorter mobile phase. Following both a short and a regular mobile phase, compaction induces a similar increase in segregation; however, due to the lower initial value, the segregation reached after a short mobile phase is lower than the one reached after a regular mobile phase. In the snapshots (Fig S3D), we show the initial and final states for systems undergoing compaction after a regular and short mobile phase.

### Effect of segregation axis alignment

As detailed in *Simulation Protocol – P2 (Compaction)*, we allow compaction only outside of a region of width *W=*0.75 *R* centered at the mid-cell. In simulations, we align this region perpendicularly to the axis of maximum segregation, which is determined using LDA. Here, we also investigated the behavior of the system during the compaction phase when the alignment of the “no-compaction” zone is instead chosen randomly. The two cases (aligned and misaligned) are compared (Fig S3E). We find that, in the misaligned case, compaction leads to an increase of segregation, but to a smaller extent than the one obtained in the aligned case. We thus conclude that compaction must take place along the axis of maximum segregation in order to reach the best possible chromosome individualization. In the snapshots in (Fig S3E), we show the initial and final states for systems undergoing fast compaction for the aligned and misaligned case.

### Effect of parameter *W* (width of “no-compaction” zone)

The parameter *W* represents the width of the mid-cell region in which compaction is inhibited (see *Simulation protocol*). Throughout this work, we have set *W=* 0.75*R*, with *R* the mean vesicle radius. In (Fig S3F), we show the extent of segregation during the compaction phase, quantified both by the segregation efficiency and the DNA-DNA distance between the two chromosomes, for *W=* 0.75*R*, 0.5*R*, and 0 (the latter corresponding to uniform compaction throughout the cell). One can see that the results remain largely unchanged for *W=* 0.5*R*, whereas for *W=* 0 the DNA-DNA distance is significantly lower, signaling a lower degree of segregation. In the snapshots in (Fig S3F), we show the initial, intermediate and final states of systems undergoing fast compaction with *W=* 0.75*R* and *W=* 0. One can see that if *W=* 0, compaction happens uniformly throughout the cells and the two chromosomes compact into a single mass, which eventually associates to the membrane.

